# A membrane-permeable small molecule biosensor accesses intractable cells and animals without genetic manipulation

**DOI:** 10.64898/2026.05.22.727289

**Authors:** Gabriel Kreider, Christopher J. MacNevin, Pothiappan Vairaprakash, Marie Rougie, Michael V. Tran, Jason J. Yi, Ewan McGhee, Denis Tsygankov, Yoshikazu Ohno, Kurt Anderson, Klaus M. Hahn

## Abstract

Fluorescent biosensors have proven valuable for revealing the spatio-temporal dynamics of protein conformation in live cells and animals. The great majority of biosensors are genetically encoded, but genetic encoding is difficult or impossible to apply in many cases, including cells or animals with poorly understood genomes, no DNA, or sensitive to manipulation. Using biosensors without genetic manipulation could greatly simplify studies in animals, expand the range of accessible organisms, and ultimately enable application in humans. Here we explore using a membrane-permeable small molecule as a fluorescent biosensor. The drug trifluoperazine, which binds only to the active conformation of calmodulin, was covalently linked to an environment-sensing merocyanine dye to create CaMero, a biosensor of calmodulin activation. Simple incubation of CaMero in the extracellular medium, or injection in the tail vein of mice, led to sensitive real time reporting of calmodulin activity. The dye underwent a 12-fold change in fluorescence intensity upon binding to activated calmodulin, revealing waves of activation in peristaltic intestine, localization and kinetics of calmodulin activation during serum stimulation in fibroblasts, and localized activation in the single-celled marine protist foraminifera.

## INTRODUCTION

Cell signaling is controlled by the subcellular localization and timing of protein conformational changes. The same proteins can induce behaviors as different as apoptosis or proliferation, depending on the spatio-temporal dynamics of activation^1,2^. This aspect of signaling is difficult to study biochemically or in fixed cells, but has been revealed by fluorescent biosensors, which have shed light on multiple aspects of cell biology. Most biosensors are genetically encoded. This provides a convenient means to introduce them into cells or animals, but can also be limiting. The need for genetic manipulation precludes studies in humans, and in cells with an intractable genome or even no genome (e.g. red blood cells, invertebrates with poorly characterized genomes, differentiated cells with low chromatin availability, and non-proliferating cells). Additionally, the creation of transgenic organisms can be challenging, expensive, and time consuming.

Here we explore a biosensor approach that can overcome these limitations. It is an extension of biosensor designs based on combining an affinity reagent (AR) with a fluorescent dye. Affinity reagents are genetically encoded polypeptides that bind selectively to the active conformation of the protein being studied. They have been computationally engineered^3,4^, or derived from fragments of downstream effectors^5-9^, antibody fragments^3,10^, and peptides^11^. In current biosensors, AR are covalently linked to dyes with solvent-sensitive fluorescence, such that fluorescence changes when the AR binds to the target protein. When loaded in cells, these biosensors have been used to report on localized conformational changes of several protein structural classes, including GTPases, kinases, growth factor receptors, and cyclin family proteins^3,5-13^.

There are substantial downsides to current biosensors based on AR. When derived from naturally occurring downstream ligands, they can compete with normal signaling interactions to create dominant negative effects. This has been overcome by using tracer amounts of biosensor with reversible binding, to rapidly sample a small proportion of the total protein pool, and exchange binding partners frequently. When possible AR binding sites are far from protein interaction interfaces, sensing activation through allosteric effects on the protein surface. Another serious disadvantage of proteinaceous AR linked to dyes has been the need for cumbersome methods such as microinjection or electroporation to load them into the cell^3,14,15^. Recently there has been work using intracellular labeling (e.g. with genetic code expansion or SNAP/HALO ligands)^12,16,17^ to generate dye-labeled biosensors in cells, but this remains challenging in many circumstances.

Here we use a membrane permeable small molecule as the AR, rather than a polypeptide. Fusing a membrane-permeable environment-sensing dye to a permeable small molecule with specific binding to the target protein’s active conformation, we generate a biosensor that is entirely a small molecule, and can enter the cell simply by diffusion from the medium (**Fig. 1a**). This can generate biosensors for the previously intractable cells and organisms described above, and could ultimately open the door to a new class of biosensors for human diagnosis, research, and surgery.

**Figure 1.**
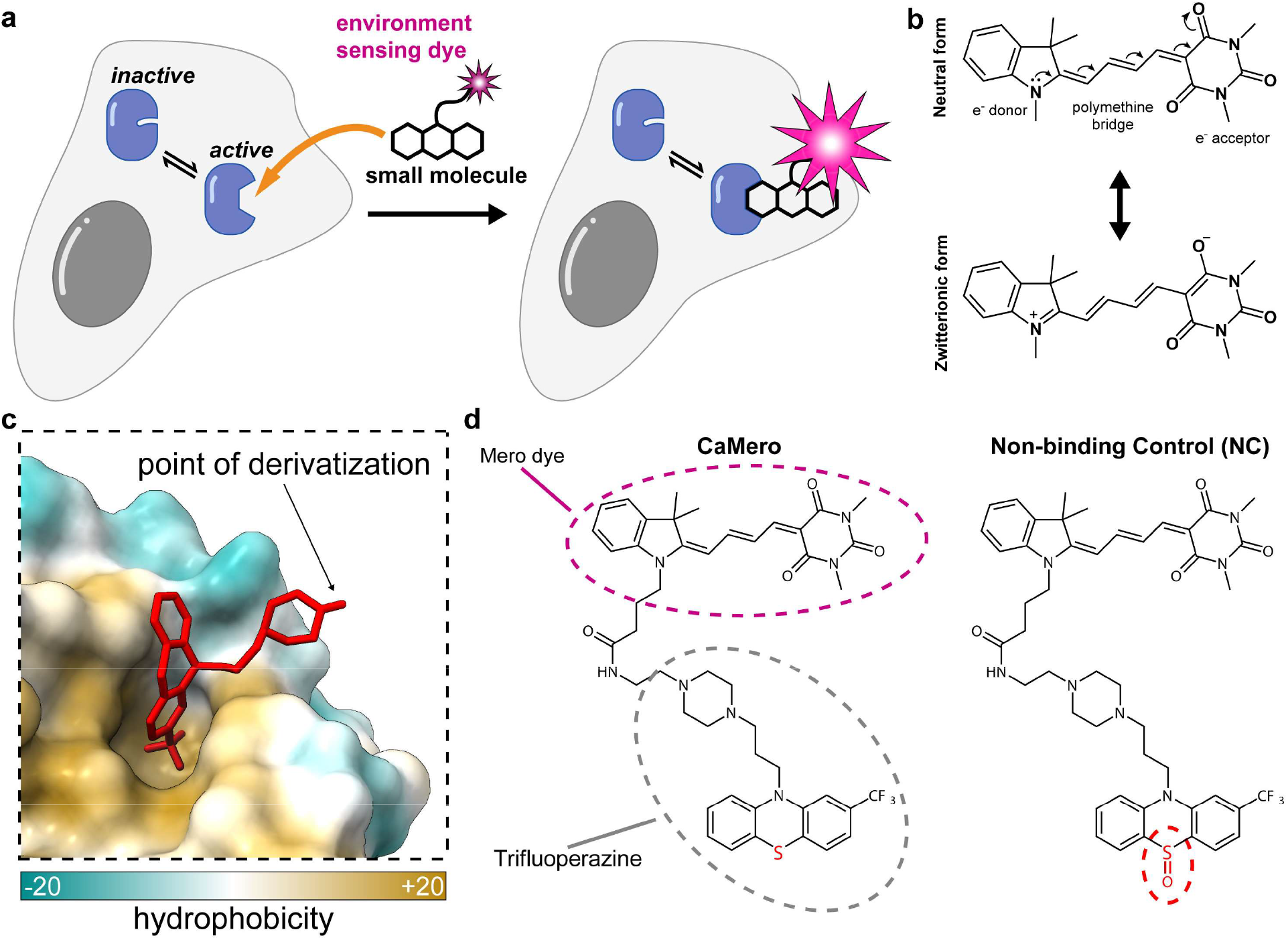
Design of CaMero. (**a**) The biosensor is a membrane permeable small molecule that enters the cell and binds specifically to the activated conformation of calmodulin. Upon binding it undergoes a 12-fold increase in fluorescence intensity. (**b**) Neutral and zwitterionic forms of the dye I-BA(4) (**c**) Crystal structure of trifluoperazine bound to calmodulin, showing the solvent exposed *N-*methyl piperazine used for dye attachment (PDB: 1LIN). (**d**) Trifluoperazine with and without sulfoxide conjugated to the environment-sensing dye I-BA(4). The sulfoxide abrogates calmodulin binding.

To test this concept, we used the calmodulin-binding molecule trifluoperazine as the AR, and attached it to dyes that were previously developed in our lab for protein-based biosensors. These environment-sensing dyes were screened for membrane permeability, and we chose dyes with bright, long-wavelength fluorescence for use in living cells^12,18,19^. The Ca^2+^-binding protein calmodulin is ubiquitously expressed in eukaryotic cells, and is involved in numerous cellular processes, including growth, motility^2^, muscle contraction^20^, and memory^2,21^. Upon binding Ca^2+^, calmodulin undergoes a conformational change to form a dumbbell-shaped structure that can interact with a range of effector proteins (**Supplemental Fig. 1**)^22^. Trifluoperazine (TFP) binds only to Ca^2+^-bound, activated calmodulin. It is an inhibitor, and so must be used at nonperturbing concentrations, but inhibitors would provide a ready source for biosensor AR. TFP also binds dopamine receptors so it is inappropriate for use in certain cells^23,24^. Ultimately, we hope to use molecules that bind allosteric sites, and so do not inhibit normal protein interactions.

We will describe the synthesis and testing of this small molecule biosensor, which we named CaMero (for calmodulin with merocyanine dye). We examine the solvent dependent fluorescence of the dye and dye-TFP conjugate. We test *in vitro* the biosensor’s interactions with calmodulin and Ca^2+^, its response to calmodulin conformational changes, and optimize the biosensors’ ability to enter mammalian cells. We use CaMero to study calmodulin activity in fibroblasts, mouse intestine, and foraminifera, a marine model organism used as a bioindicator of coastal ecosystems, ocean acidification, and as a proxy for coral reef health^25-27^.

## RESULTS

### Design and synthesis of the biosensor

We first generated a biosensor by linking an environment-sensing dye to trifluoperazine (TFP). The dye had to be attached to TFP where it would not impact the affinity and selectivity of the drug for calmodulin, but where its spectrum would be affected when TFP interacted with calmodulin. We and others have extensively characterized merocyanine dyes for use in live cell biosensors^12,19,28-30^. Merocyanines are “push-pull” dyes, composed of electron donor and electron acceptor groups linked by a polymethine bridge (**Fig. 1b**). These fluorophores can be seen as adopting a neutral or zwitterionic form, depending on the polarity and hydrogen bonding of interacting solvent molecules or protein residues^31-34^. Our laboratory previously screened a range of donor and acceptor combinations to increase solvent effects on fluorescence intensity and/or excitation/emission maxima, and to optimize brightness, photostability, and fluorescence at long wavelengths for use in live cells and animals^18,19,28,35^. Based on these studies, we chose a merocyanine dye with an indolenine donor and a barbituric acid acceptor connected by a four-carbon methine bridge, hereafter referred to as I-BA(4).

For the affinity reagent (AR) of the biosensor we used trifluoperazine, a small molecule known to bind selectively to calmodulin’s active, Ca^2+^-bound conformation^36^ (see introduction). Published crystal structures of the TFP-calmodulin complex^37-39^ showed that the piperazinyl-methyl group of TFP is not involved in calmodulin binding (**Fig. 1c**), and is exposed to the surrounding water. We therefore attached the dye at this position via a 4-carbon chain containing an amide (see supplemental methods for synthesis). Because the dye was not highly water soluble, we reasoned that it would interact with hydrophobic surfaces on calmodulin, resulting in a spectral change. The resulting biosensor was named CaMero (**Fig. 1d**). Previous studies had shown that a single atom change in TFP—adding an oxygen to the sulfur of the phenothiazine ring—greatly reduced TFP affinity for calmodulin^36,40^. The addition of oxygen was used to make a negative control biosensor, named CaMero-NC (non-binding control) (**Fig. 1d**).

We also made biosensor variants in which the donor ring of the dye bore acetoxymethyl (AM) ester groups. These have long been used to facilitate cell entry of small molecules. AM ester groups facilitate passage through the membrane by replacing charged carboxylic acids with more hydrophobic esters. Once in the cell they increase retention because intracellular esterases cleave them back to charged carboxylic acids^41-44^. These acetoxymethyl ester biosensors were named CaMero-AM and CaMero-AM-NC (see supplemental methods for structure and synthesis).

### Photophysical characterization of free dye, CaMero, and CaMero-NC

We next characterized the photophysical properties of CaMero. **Supplemental Figures 2a-j** show the absorbance and emission characteristics of the unfunctionalized dye, and the dye connected to TFP. These compounds were examined in solvents of varying polarity and hydrogen bonding capacity. The strongest emission was produced in solvents that were both hydrogen bond donors and nonpolar (1-octanol, t-butanol). Intermediate brightness was seen with methanol, a hydrogen bond donor but relatively polar, and chloroform, which is nonpolar but only a weak hydrogen bond donor. Nonpolar hydrogen bond acceptors (dioxane and THF) showed low brightness values, except for the more polar DMSO, which produced bright fluorescence. This is consistent with previous molecular modeling studies indicating that hydrogen bond donors interacting with the terminal oxygen of merocyanine acceptors play an important role^32-34^. Thus, the dye was likely to respond to calmodulin when it encountered a hydrophobic environment on the protein’s surface, and could interact with side chains via specific hydrogen bonds. The dye was capable of strong fluorescence emission—the extinction coefficient times quantum yield exceeded that of Alexa Fluor 555, Cy3, ATTO 647 and approached Alexa Fluor 750 and ATTO 700, dyes commonly used in protein imaging studies (Alexa Fluor 555: 15,000, Cy3: 22,500, ATTO 647: 24,000, CaMero: 28,600, Alexa Fluor 750: 28,800, and ATTO 700: 30,000). Adding TFP to the fluorophore enhanced the dye’s brightness slightly, while maintaining the same trends in solvent-dependence (**Supplemental Fig. 2d-j**).

The absorption spectra of the biosensors CaMero and CaMero-NC were recorded in aqueous Tris buffer and phosphate buffered saline (**Supplemental Fig. 3a-f**). Based on previous studies of merocyanine dyes^45,46^ the spectra indicated that CaMero might aggregate in aqueous solution (**Supplemental Fig. 3a**). This was further tested by measuring changes in emission upon dilution, assuming that aggregation would be concentration-dependent (**Supplemental Fig. 3g and h**). The molar emission of CaMero-NC remained stable while CaMero’s emission increased upon dilution, indicating that CaMero was aggregating at higher concentrations. The biosensor likely did not pass into the cell as an aggregate, or aggregate appreciably at the concentrations in the cell, but if this occurred it would reduce the brightness of unbound biosensor, enhancing sensitivity. Aggregation could also reduce fluorescence background in the medium. The studies below show that there was sufficient interaction of the biosensor with calmodulin, whatever its form.

### Biosensor response to calmodulin *in vitro*

Using aqueous Tris buffer, we examined the response of the AM-containing biosensors to calmodulin and Ca^2+^. In the presence of excess calmodulin, CaMero-AM showed an 8-fold increase in fluorescence intensity when Ca^2+^ concentration was increased (**Fig. 2a**). Minimal change was observed in the absence of calmodulin, and the negative control probe CaMero-AM-NC showed only a very weak response, likely due to low, residual affinity for calmodulin.

**Figure 2.**
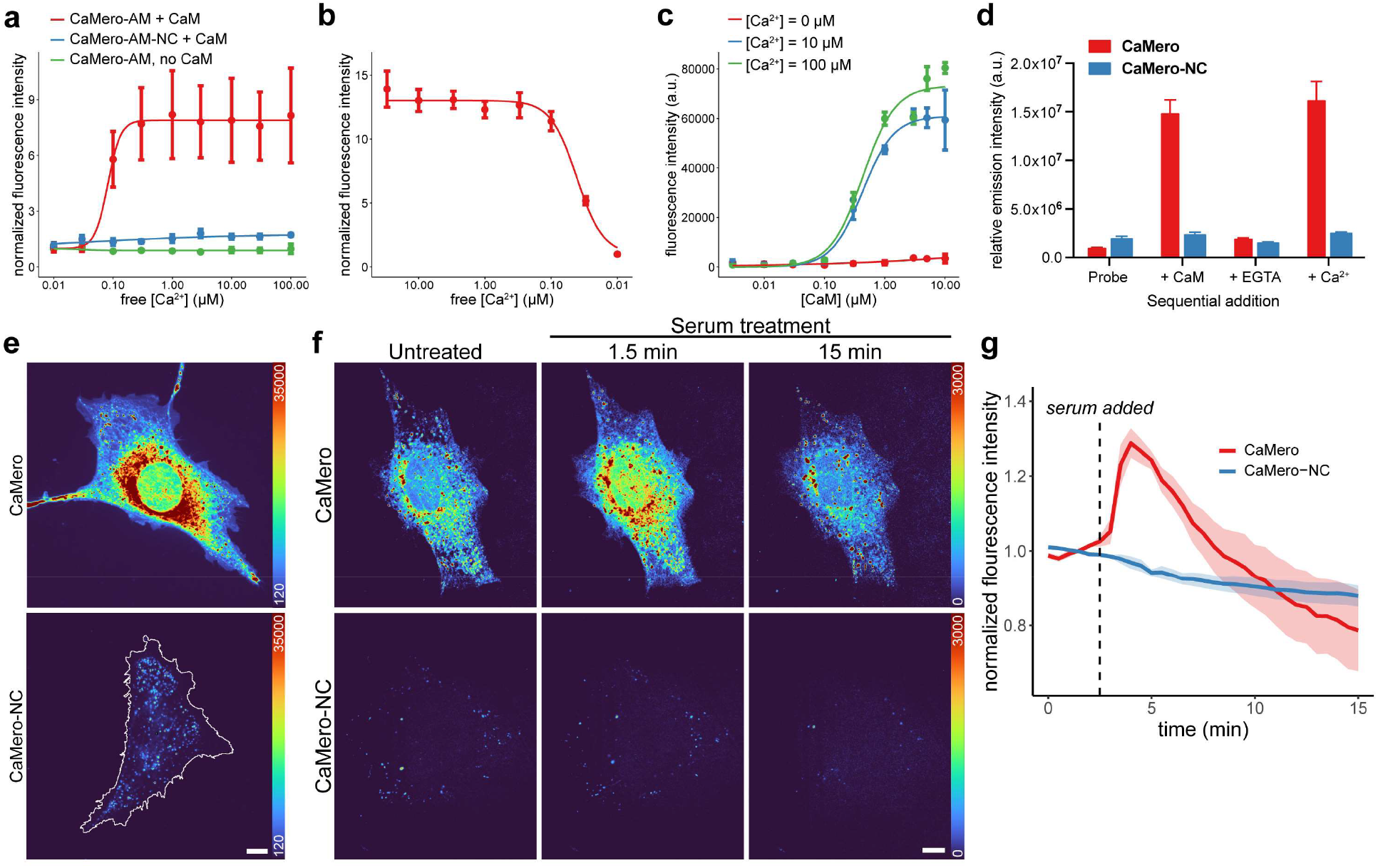
CaMero changes fluorescence only in the presence of both calcium and calmodulin. (**a**) CaMero-AM and control CaMero-AM-NC in calmodulin solution with increasing Ca^2+^. **b**) calmodulin-induced fluorescence increases are reversible. (**c**) Increased CaMero-AM fluorescence induced by Ca^2+^/calmodulin is reversed by addition of EGTA. (**d**) Emission of CaMero and CaMero-NC during sequential addition of calmodulin, EGTA and Ca^2+^. (**e**) MEF cells incubated with CaMero show significantly greater fluorescence intensity than CaMero-NC. Images were scaled identically, with linear scale selected to show range of CaMero intensities across the cell. (10-micron scale bar). (**f**) MEF cells incubated with CaMero or CaMero-NC probe were serum starved and then treated with FBS. Images were scaled identically, with scale selected to prevent camera saturation at peak CaMero intensity (10micron scale bar). (**g**) Fluorescence intensity change upon serum stimulation for CaMero and CaMero-NC control (imaged every 15 s for 15 min, FBS added at 3 min, mean ± s.e.m. of 3 replicates with at least 15 cells per condition).

We tested reversibility by adding ethylene glycol tetraacetic acid (EGTA) to a solution of calmodulin, Ca^2+^, and biosensor. This produced a drop in fluorescence, occurring between 0.1 and 0.01 μM free Ca^2+^ (**Fig. 2b**). To examine the K_d_ and cooperativity of biosensor-calmodulin binding, we titrated the Ca^2+^ concentrations of solutions containing calmodulin and biosensors (**Fig. 2c**). The resulting Hill curves showed a sigmoidal response to calmodulin with a Hill coefficient of 1.68 ± 0.22, indicating cooperative binding of more than one CaMero-AM molecule to calmodulin, and a K_d_ of 0.43 μM (values for 10 μM free Ca^2+^). It has been established that calmodulin has multiple sites that can bind TFP with different affinity^36-38^. The K_d_ we observed was close to the reported value of 0.95 μM^40^.

When testing CaMero without acetoxymethyl esters, we added excess calmodulin (60.0 μM) to a solution of CaMero (1.00 μM) in aqueous Tris buffer containing 10.0 μM Ca^2+^. This induced a 12-fold increase in fluorescence, which could be brought back to the initial fluorescence level by adding EGTA. Adding Ca^2+^ to this solution restored the maximum fluorescence intensity (**Fig. 2d, Supplemental Fig. 4**), demonstrating reversibility. We twice reversed the fluorescence response of the CaMero without AM esters by first adding calmodulin, then Ca^2+^, and then EGTA. The control biosensor CaMero-NC showed minimal response.

Together, these studies demonstrated that interaction of the biosensor with calmodulin produced a large fluorescence increase, was reversible, and was induced by Ca^2+^-bound calmodulin and not by Ca^2+^ or calmodulin alone.

### Spatio-temporal dynamics of calmodulin activation in live cells

With *in vitro* characterization in hand, we set out to use the biosensors in living cells. We chose NIH 3T3 mouse embryonic fibroblast cells (MEFs) because we are well familiar with their behavior, responses to other fluorescent probes, and signs of ill health. We simply incubated the biosensor in the extracellular medium and in some experiments washed them to remove extracellular biosensor. Different incubation times and concentrations were tested, monitoring the increase in intracellular fluorescence, and the coincident production of fluorescent vesicles due to endocytic uptake. Good calmodulin labeling with minimal vesicles was achieved by incubating in 0.5-1.0 μM biosensor for 20 minutes, followed by 3 washes (see Methods). Washing was not necessary for all experiments, because the biosensors produced minimal fluorescence when they were not bound to calmodulin. The CaMero and CaMero-AM variants were compared in cells with and without Pluronic F-127 (PF), which is commonly used to enhance solubility and permeability with AM esters^47^. CaMero without the AM groups and without PF produced the greatest fluorescence intensity (**Supplemental Fig. 5a and b**), so these conditions were used in most of the experiments below. To assess the ability of CaMero without AM esters to stay available within the cell, cells were examined before and after washing (Methods, **Supplemental Fig. 5c**). Washing caused a gradual decrease of intracellular biosensor over 2 hours. Longer experiments may require the use of low concentrations of probe in the media, or use of the AM ester form of the biosensor.

When serum-starved cells loaded with biosensor were stimulated with serum, they showed a rapid 30% increase in calmodulin activity, peaking within 5 minutes and then more slowly dropping off (**Fig. 2f, g, and Supplementary Video 1**). Thus, the new biosensor correctly reported well known calmodulin activation kinetics in response to the release of Ca^2+^ from intracellular stores^48-51^. The control CaMero-NC biosensor showed no response to serum addition.

Although cells incubated in biosensor showed fluorescence throughout the cell, the great majority of fluorescence was concentrated in the perinuclear region (**Fig 2e**). This was consistent with previous calmodulin biosensor studies^48^, and notably different from the distribution of GFP-tagged calmodulin^52^, or Ca^2+^ as shown by live cell Ca^2+^ biosensors^16,53^. Thus, calmodulin does not simply transmit changes in Ca^2+^ dynamics to downstream molecules, but further localizes those downstream effects. CaMero-NC, the control biosensor, showed only weak fluorescence.

I-BA(4) showed solvent-dependent changes in intensity, but minimal shifts in excitation or emission maxima. Intensity variations were used above to monitor changes in the overall calmodulin activity of individual cells, and to visualize the localization of calmodulin activity, but precise quantitation of intensity measurements is susceptible to artifacts from subcellular variations in biosensor concentration or potentially by non-specific interactions of dyes^54,55^.

These artifacts can be overcome by using fluorescence lifetime imaging (FLIM)^55^, or by introducing a second, activity-insensitive fluorophore for normalization. We imaged CaMero and CaMero-NC using fluorescence lifetime microscopy and analyzed the results using phasor plots and Gaussian mixture model (GMM) cluster analysis (**Supplemental Fig. 6a and b**)^56,57^. The CaMero probe had a fluorescence lifetime around 1.2 ns while the CaMero-NC probe had a longer fluorescence lifetime of around 1.7 ns. GMM clustering of the CaMero phasor plot was used to separate distinct clusters that contained either the desired signal (cluster 1) or vesicular biosensor (cluster 2). CaMero-NC was found to be primarily in cluster 2. Since the biosensor bound to calmodulin has a distinct lifetime, other fluorescent signals could be eliminated. This could be valuable in cells (e.g. vesiculation obscures simple intensity signals) or in other cases where unbound CaMero needs to be distinguished from bound.

### Calmodulin activation in challenging organisms

An important advantage of a membrane-permeable small molecule biosensor is its ability to be used in situations where manipulation of the genome is cumbersome or impossible. We tested whether CaMero could be introduced into a mouse by examining calmodulin dynamics in peristaltic intestine, which undergoes periodic contractions known to be driven by Ca^2+^/calmodulin^58^. CaMero-AM or CaMero-AM-NC were injected into GFP-LifeAct mice and sections of intestine were imaged ex vivo using multi-photon microscopy (**Fig. 3a**). Intestines from CaMero-AM mice were substantially brighter than those from mice loaded with CaMero-AM-NC. Image stacks were acquired starting at the base of the crypt of Lieberkühn (Z = 0 µm) downwards through the circular (Z = -56 µm) and longitudinal (-84 µm) smooth muscle layers, which were visualized using GFP-LifeAct. CaMero-AM was localized in the fibers of each muscle layer, while CaMero-AM-NC did not clearly localize to smooth muscle cells. Widefield imaging of intestine sections also showed bright signal with CaMero-AM while CaMero-AM-NC was much weaker (**Fig. 3b and c**). The excised tissue maintained contractility ex vivo, and quantification of fluorescence intensity in the tissue sections showed spikes in fluorescence intensity with each contraction (**Fig. 3d**). No such periodic oscillations were observed with CaMero-AM-NC.

**Figure 3.**
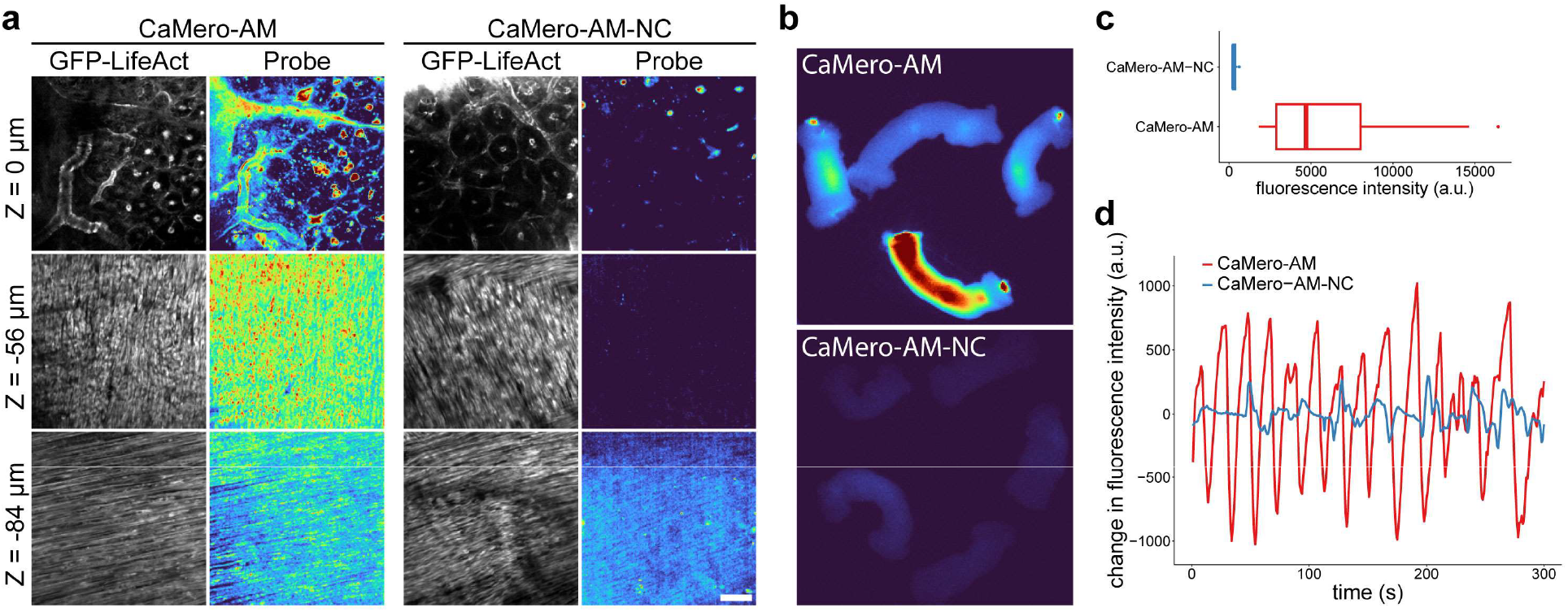
GFP-LifeAct mice injected with biosensors, 2 photon imaging. (**a**) Two-photon imaging of smooth muscle layers in excised intestine from mice injected with biosensor (CaMero-AM or nonresponsive control CaMero-AM-NC; 50-micron scale bar). (**b**) Widefield fluorescence microscopy of excised intestine shows intensity of CaMero-AM relative to CaMero-AM-NC. (**c**) Quantification of panel (**b**) for at least 3 mice per condition. (**d**) Change in fluorescence intensity over time for CaMero versus CaMero-NC. The CaMero probe shows periodic changes in intensity consistent with the frequency of tissue contraction.

Finally, we tested CaMero in foraminifera, a marine model organism frequently used as a bioindicator of coral reef health (**Fig. 4a**)^25-27^. Foraminifera have calcium carbonate shells called tests, with calmodulin highly expressed during formation^25^. Fine pseudopodia, called reticulopodia, extend from the tests and function in locomotion, attachment, and food gathering^59-62^. After treatment, CaMero signals were enriched in the filamentous reticulopodia protruding from the tests (**Fig. 4b**), whereas the non-binding control CaMero-NC produced little signal. Unlike calcein-AM^61^ or SiR-actin^63^, CaMero did not simply label cytoplasmic or cytoskeletal structures, but reported localized calmodulin activity in reticulopodia. When we added EGTA-AM to chelate intracellular Ca^2+ 64^, calmodulin activity was reduced (**Fig. 4c and Supplementary Video 2**). This was reversed by adding seawater to restore normal Ca^2+^. During recovery from EGTA-AM treatment, the reticulopodia associated with foraminifera motility were also restored (**Fig. 4d**). Mechanical stress on reticulopodia during foraminifera translocation was associated with localized spikes in calmodulin activity (**Fig. 4e**).

**Figure 4.**
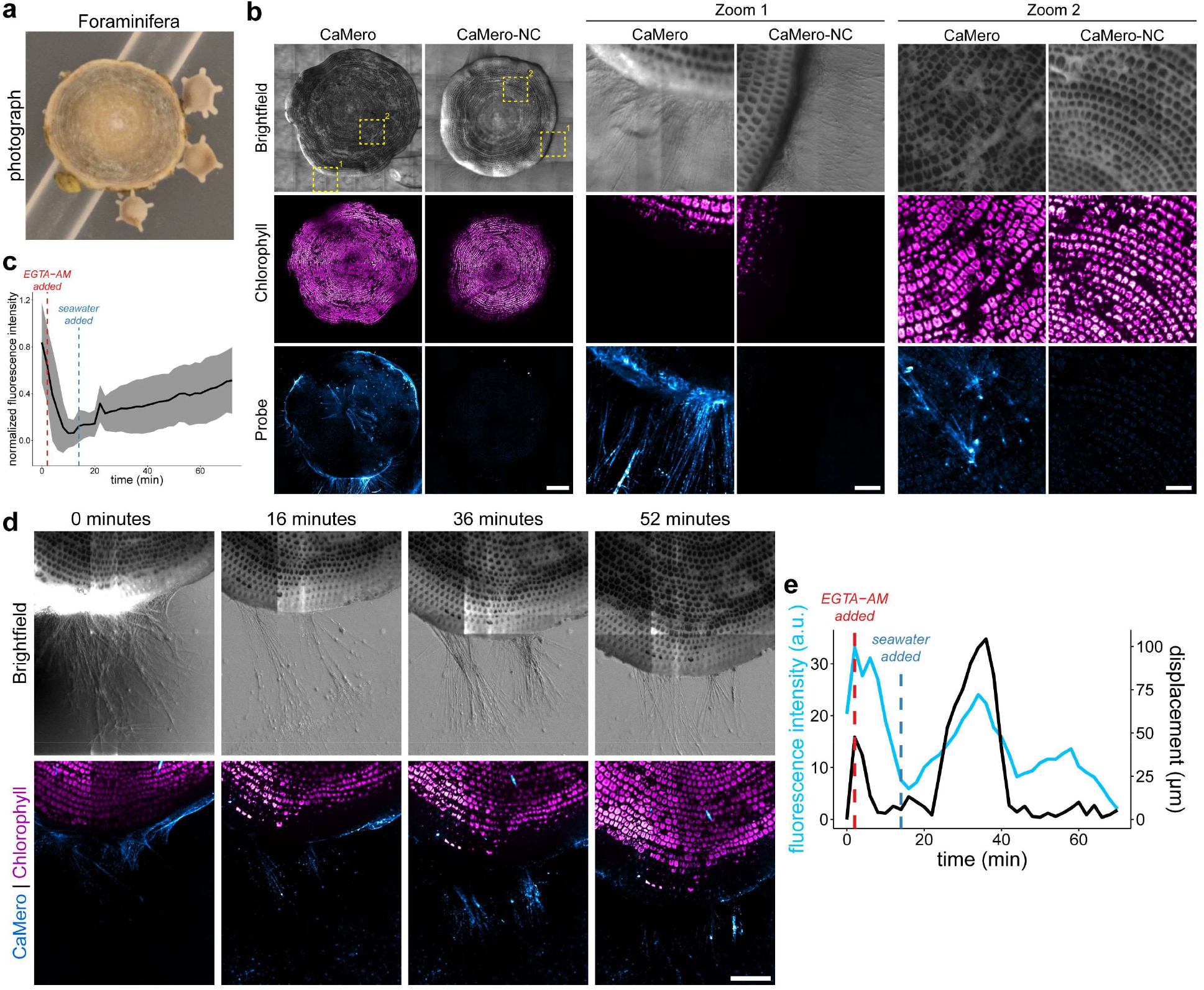
CaMero shows calmodulin activity in the reticulopodia of foraminifera. (**a**) Photograph of an Amphisorus kudakajimensis surrounded by three Calcarina gaudichaudii large benthic foraminifera, sea-dwelling eukaryotes that form a calcareous shell (largest approximately 0.5 cm in diameter). (**b**) CaMero or CaMero-NC in foraminifera (Amphisorus kudakajimensis). Chlorophyll fluorescence is from symbiotic algae (zooxantela) within the cytoplasm (1 mm scale bar for whole cell; 100 μm for zoom). (**c**) Loss of CaMero fluorescence upon treatment with EGTA-AM. Recovery after adding seawater. (**d**) A foraminifera moving along the coverslip after recovery from EGTA-AM treatment. The field of view remains fixed to show movement towards the bottom of the frame. Flashes of calmodulin activity in reticulopodia during translocation (1 mm scale bar). (**e**) Fluorescence intensity over time from experiment in panel **d**.

## DISCUSSION

We have demonstrated that a fluorescent biosensor consisting simply of a small molecule can produce a bright, specific signal to report protein conformation, in cultured cells, mice, and marine invertebrates. This provides a substantial advance in convenience—simply add the biosensor to the medium, wash only if necessary, and begin experiments. CaMero was used to image protein activity in mice without the substantial effort required to make a transgenic animal, and enabled ready multiplexing with an already expressed fluorescent protein. The foraminifera we studied were unsuitable for genetic techniques due to their sensitivity and lack of available genetic tools.

CaMero was based on a known calmodulin inhibitor, even though inhibitors can perturb cell behavior. This highlights an important limitation. There are few known small molecules that bind to a specific protein conformation and minimally inhibit protein interactions or activity. Although drugs and known inhibitors were selected because they affect protein activity, they do provide a large pool of AR candidates, and previous work has shown that inhibitory proteins can be used in biosensors. Published AR have often been based on fragments from proteins downstream of the target, because these effectively interact only with the active conformation^5-8^. Dominant negative effects were overcome by using tracer amounts of AR with K_d_ weakened for reversible interaction, but still strong enough to maintain specificity^3,15,65^. More recently, there has been impressive progress in identifying small molecules computationally^66-70^. Some drugs bind to conformationally-dependent allosteric sites at regions far from ligand interaction surfaces^71,72^. These have been designed computationally, and could be selected for minimal effects on protein activity. It will be important to leave interactions with upstream ligands intact, as these generate the very changes in activity that biosensors are meant to monitor. For imaging, one must also maintain the interactions that cause localization. Methods to test and optimize this have been described^5,7,15,73^.

Like biosensors based on fluorescent substrates^13,74-76^, the AR-dye design we use here reports on endogenous proteins, potentially reducing cell perturbation. Unlike substrates, it cannot drift away after its fluorescence has been altered, so can enhance spatial resolution. CaMero benefits from using an environment-sensing dye rather than fluorescence resonance energy transfer (FRET). The dye is directly excited rather than indirectly via energy transfer, so can be substantially brighter. The limitation of sensing endogenous protein is the potential for off-target binding. For example, TFP is known to block dopamine receptors^23,24^ so CaMero may not be appropriate for use in cells or tissues with high dopamine receptor expression.

TFP presented an obvious position where the dye could be attached without interfering with TFP-calmodulin binding. Despite having no high affinity interaction with calmodulin, the dye at that position underwent a 12-fold change in fluorescence intensity when it was brought near calmodulin by TFP. This was likely because the dye preferred to interact with a hydrophobic region on calmodulin rather than water. Such a surface might also provide hydrogen bonds to the carbonyl groups on the dye, a position known to produce fluorescence changes^32-34^. Affinity for hydrophobic surfaces can be valuable, but could also lead to nonspecific interactions. Our previous work indicates that some dyes tend to stain lipid bound organelles including endoplasmic reticulum and vesicles^12,28^. I-BA(4) proved to be a useful dye that did not show appreciable nonspecific staining or endocytic uptake, but produced a large fluorescence change in the CaMero biosensor. The structure of the dye can impact whether it undergoes a fluorescence response for any given protein, regardless of binding affinity. Molecular modeling and experimental studies have shown that hydrogen bonds to specific heteroatoms within dyes can produce strong fluorescence changes^32-34^. A toolchest of dyes with different distributions of charged and hydrophobic regions will be useful. Some dyes have a tendency to undergo constitutive endocytosis^77^. The tradeoff between passage through the cell membrane and endocytic uptake depends on cell type and biosensor, and will likely require optimization of incubation time and concentration to obtain sufficient signal strength while avoiding bright vesicles. FLIM imaging could be useful in such cases (**Supplemental Fig. 6a and b**).

In MEFs, we observed high calmodulin activity near the nucleus. This was consistent with previous studies^48^, including localization of active calmodulin at internal membranes and channels in the endoplasmic reticulum^2,78^. The active calmodulin rapidly and transiently increased in response to serum stimulation, likely due to a rapid influx of Ca^2+^ and/or release from internal stores^48,79^. In smooth muscle contraction, calmodulin activates myosin light-chain kinase, which phosphorylates myosin^80^. We were able to visualize endogenous calmodulin activity in the layers of intestinal smooth muscle cells within two hours of injecting CaMero. CaMero provided a signal bright enough to observe changes in fluorescence intensity at low magnification in peristaltic tissue sections. Finally, our studies in foraminifera demonstrated that small molecule biosensors could be applied to study signaling in challenging organisms that do not have available genetic tools. We were able to observe high calmodulin activity in the reticulopodia and saw spikes in activity associated with tension during translocation. Although calmodulin-related molecular changes have been reported in foraminifera^25^, we are not aware of any previous work that has directly visualized calmodulin activity in living foraminifera.

We hope that this example using a dye-small molecule combination as a biosensor will stimulate application of biosensors to previously intractable systems. Ultimately small molecule biosensors could be applied to humans, for rapid diagnosis, surgery, or to monitor intracellular protein activities with a simple blood test. Although applications using existing drugs are possible, small molecule biosensors can build upon new methods in computational drug discovery to identify binders of allosteric sites. The future will likely bring dyes that undergo ratiometric fluorescence changes, dyes whose brightness will enable reduced biosensor concentration, and dyes in multiple wavelengths for multiplexed imaging.

## METHODS

### *In vitro* assays

To maintain specific Ca^2+^ concentrations, stock Ca^2+^ solutions were prepared with 100 mM NaCl and 50 mM Tris-HCl, pH 7.5. These were buffered with the appropriate amount of EGTA to maintain a targeted free Ca^2+^ concentration at an ionic strength of 0.15, as described^64^. Amounts of EGTA were determined using the online calculator: https://somapp.ucdmc.ucdavis.edu/pharmacology/bers/maxchelator/CaEGTA-TS.htm^81^. Calmodulin was purchased from Sigma-Aldrich (cat. P1431). Fluorescence intensity experiments were run in triplicate in 386-well plates. For Ca^2+^ titration and reversibility experiments, calmodulin and biosensor (CaMero or CaMero-NC) were both 1.0 µM.

### Cell culture and treatment with CaMero biosensors

NIH 3T3 mouse embryonic fibroblasts (ATCC) were maintained in 5% CO_2_ at 37 C° in Dulbecco’s modified Eagle’s medium (DMEM, Cellgro) with 10% fetal bovine serum (HyClone, Thermo Scientific) and 2 mM GlutaMAX (Gibco, Life Technologies). Cells were plated 3-4 hours before imaging on coverslips coated with fibronectin 20 μg/mL (Sigma-Aldrich) overnight. Cells were incubated with biosensor for 20 min in 5% CO_2_ at 37 °C in imaging medium [FluoroBrite DMEM (A1896701, Gibco, Life Technologies) with 5% fetal bovine serum (HyClone, Thermo Scientific) and 1X GlutaMax (Gibco, Life Technologies)]. Titration of CaMero and CaMero-NC in MEFs determined the optimal concentration of CaMero for cell culture to be around 0.5-1.0 μM. CaMero intensity plateaued around 1 μM and the negative control remained consistently low across the range of concentrations (**Supplemental Fig. 7**). The cells were gently washed with imaging medium (3 × 1 mL) and imaged in the same medium.

### Confocal imaging

Live cell imaging was performed on a custom spinning disk confocal microscope based on an IX81 with a ZDC2 autofocus system. The microscope was equipped with a CSU-X1 spinning disk (Yokogawa), 6 solid-state lasers (Coherent OBIS Galaxy 405, 445, 488, 514, 561, 640), two dichroic mirrors (AND-0007:405/491/560-570/647 and AND-0005:457/515/647, Andor), seven emission filters (FF01-446/523/600/677, FF01-483/32, FF01-525/30, FF01-542/27, FF01-607/36, FF02-685/40, FF02-447/60, Semrock) in a 10-position filter wheel (Sutter Instruments), image splitting optics (TuCam, Andor), and an XYZ stage with piezo Z-axis (Applied Scientific Instrumentation). Fluorescence images were collected using either a 60X objective (UPLAPO60XOHR, NA 1.5, Olympus) for MEF imaging or a 4x objective (UPLFLN4XPH, NA 0.13, Olympus) for foraminifera imaging. Images were obtained using a scientific complementary metal oxide semiconductor (sCMOS) camera (Fusion BT, Hamamatsu). MEF cells were imaged in a stage top incubator with 5% CO_2_ and humidity (Uno, Okolab). The system was controlled by the open-source software Micromanager^82^. All images where processed for dark current correction, shade correction, background subtraction, and photobleaching correction as applicable in FIJI^83^.

### FLIM imaging

All FLIM experiments were performed on a Leica STELLARIS 8 FALCON STED microscope with a 63X objective (Plan-Apo, NA 1.4, Leica). The microscope was equipped with a white-light laser (440 nm - 790 nm) plus a 405 nm diode laser, 3 HyD S detectors and 2 HyD X detectors, AFC (Automatic Focus Correction), and a stage top incubator (Tokai Hit). FLIM phasor analysis was performed using GSLab^57^.

### Serum stimulation experiments

Glass-bottom dishes (MatTek) were coated with 5 μg/mL fibronectin in PBS, incubated for at least 30 minutes at 37 C°, and washed 3X with PBS. MEFs were plated in DMEM with 10% FBS and 1X GlutaMAX (Gibco, Life Technologies). The following day, cells were washed with PBS 3X, and the medium was replaced with starvation medium overnight (0.5% FBS in FluoroBrite DMEM with 1X GlutaMAX). CaMero and CaMero-NC were then diluted to 100 nM in warmed starvation imaging media. Cells were incubated in imaging medium with dye for 20 minutes in a 5% CO_2_ incubator at 37 C° before 3X washing with pre-warmed PBS. 2 mL starvation medium was added to each dish, and the cells were placed back in the CO_2_ incubator for 10 minutes prior to imaging. The cells were then imaged every 30 seconds with 100 ms exposure using a 561 nm laser set to 74 µW at the objective. After 3 minutes of imaging, the cells were stimulated with 90 μL FBS to achieve a final concentration of 5% FBS.

### Mouse intestine experiments

Treatment solutions of the probes were prepared as follows: 8.0 µL stock solution of probe (CaMero-AM or CaMero-NC-AM at 20 mM in DMSO), 8.0 µL 20% w/v Pluronic F-127 in DMSO, 15 µL stock solution of Hoechst 33342 (10mM in DMSO), and 120 µL phosphate buffered saline were combined and mixed in the order listed to give a final concentration for the probe of approx. 1.0 µM and a total working volume of 150 µL. Treatment solutions were administered to GFP-LifeAct mice via tail-vein injection. Mice were allowed to recover for a period of 1-2 hours, sacrificed by cervical dislocation, and short sections (∼1 cm) of the intestine were removed at approximately 10 - 15 cm from the stomach, washed once in PBS, and placed into Fish Ringer’s solution in glass bottom dishes (MatTek) for imaging. Tissue was imaged using a LaVision TRIM scope attached to a Nikon Eclipse Ti inverted stand with an Olympus 20x, 0.95 NA water immersion objective. Two photon excitation of CaMero-AM/CaMero-AM-NC was achieved with an optical parametric oscillator laser tuned to 1110 nm and emission was detected using a 630/60 emission filter for CaMero-AM/CaMero-AM-NC and a 549/15 filter for the SHG signal of collagen. Hoechst was excited with a Ti-sapphire laser tuned to 800 nm, and emission was collected using a 435/40 filter. For localization studies, tissue samples were imaged from the crypt of Lieberkühn and extending through to the peripheral smooth muscle layer using incremental steps in the Z plane of 4 µM (30 frames in total). CaMero-AM and GFP-LifeAct images were displayed using different contrast settings for each axial position. The same contrast settings were applied to images of the CaMero-AM and CaMero-AM-NC probes. For imaging of muscle contraction, intestine sections were imaged using an Olympus OV-100 wide-field epi-fluorescence microscope. CaMero-AM and CaMero-AM-NC samples were acquired within approximately 10 minutes of each other using identical acquisition parameters.

### Foraminifera experiments

Foraminifera were collected from Okinawa, Japan and maintained on the bench under regular room lights at room temperature. Foraminifera were maintained in filtered natural seawater collected from the Ogasawara Islands (0.2 µm; Advantec Co., Ltd., Japan). Seawater was replaced every week, and experiments were conducted within 2-3 weeks of collecting. Only samples that had visible reticulopodia were imaged. Foraminifera were transferred to glass-bottom dishes (MatTek) and allowed to attach for at least one hour. Seawater containing 0.5 μM CaMero or CaMero-NC was added to the samples and incubated for 30 minutes before being washed 3x with seawater. After the last wash, 500 μL seawater containing 50 nM of CaMero was used to maintain biosensor availability over the course of imaging. EGTA-AM treatments were done with 1 mM EGTA-AM in seawater and treated for the indicated times. 5 mL of seawater were added after EGTA-AM treatment to restore intracellular Ca^2+^ levels.

### Additional Methods

Synthetic details and characterization data for compounds I-BA(4), CaMero, CaMero-NC, CaMero-AM, and CaMero-AM-NC are described in the **Supplementary Methods**.

## Supporting information

Supplementary Video 1. Calmodulin activity reported by CaMero in MEF cells responding to serum stimulation

Supplementary Video 2. Calmodulin activity in a foraminifera treated with EGTA-AM and seawater

## ACKNOWLEDGMENTS

We thank Gary L. Johnson for valuable discussions. We gratefully acknowledge the American Cancer Society (C.J.M, 119169-PF-10-183-01-TBE), the National Institutes of Health (R35-GM122596, KMH), and the Japan Science and Technology Agency (PRESTO, JPMJPR24G2, YO) for financial support. We are grateful to the UNC Lineberger Comprehensive Cancer Center and the UNC Hooker Imaging Core for providing the Leica STELLARIS for FLIM imaging (P30 CA016086 Cancer Center Core Support Grant; NIH 1S10OD030300). UNC core facilities supporting this work were funded by the National Science Foundation (grant no. CHE1726291 for the Mass Spec facility), and the NIH (S10OD032476 for upgrading the 500 MHz NMR spectrometer).

## Notes

### Competing Interest Statement

The authors have declared no competing interest.

### Summary of Updates

Additional references included. Minor text edits, formatting, and figure revisions.

## REFERENCES

1 Jaffe, A. B. & Hall, A. Rho GTPases: biochemistry and biology. Annu Rev Cell Dev Biol 21, 247–269 (2005). 10.1146/annurev.cellbio.21.020604.150721

2 Chin, D. & Means, A. R. Calmodulin: a prototypical calcium sensor. Trends in Cell Biology 10, 322–328 (2000). 10.1016/S0962-8924(00)01800-6

3 Gulyani, A. et al. A biosensor generated via high-throughput screening quantifies cell edge Src dynamics. Nat Chem Biol 7, 437–444 (2011). 10.1038/nchembio.585

4 Kummer, L. et al. Knowledge-Based Design of a Biosensor to Quantify Localized ERK Activation in Living Cells. Chemistry & Biology 20, 847–856 (2013). 10.1016/j.chembiol.2013.04.016

5 Pertz, O., Hodgson, L., Klemke, R. L. & Hahn, K. M. Spatiotemporal dynamics of RhoA activity in migrating cells. Nature 440, 1069–1072 (2006). 10.1038/nature04665

6 Hodgson, L., Pertz, O. & Hahn, K. M. in Methods in Cell Biology Vol. 85 63–81 (Academic Press, 2008).

7 Kraynov, V. S. et al. Localized Rac activation dynamics visualized in living cells. Science 290, 333–337 (2000). 10.1126/science.290.5490.333

8 Mahlandt, E. K., Kreider-Letterman, G., Chertkova, A. O., Garcia-Mata, R. & Goedhart, J. Cell-based optimization and characterization of genetically encoded location-based biosensors for Cdc42 or Rac activity. J Cell Sci 136 (2023). 10.1242/jcs.260802

9 Itoh, R. E. et al. Activation of rac and cdc42 video imaged by fluorescent resonance energy transfer-based single-molecule probes in the membrane of living cells. Mol Cell Biol 22, 6582–6591 (2002). 10.1128/mcb.22.18.6582-6591.2002

10 Szent-Gyorgyi, C. et al. Fluorogen-activating single-chain antibodies for imaging cell surface proteins. Nature Biotechnology 26, 235–240 (2008). 10.1038/nbt1368

11 Liu, Q., Wang, J. & Boyd, B. J. Peptide-based biosensors. Talanta 136, 114–127 (2015). 10.1016/j.talanta.2014.12.020

12 MacNevin, C. J. et al. Membrane-Permeant, Environment-Sensitive Dyes Generate Biosensors within Living Cells. Journal of the American Chemical Society 141, 7275–7282 (2019). 10.1021/jacs.8b09841

13 Frei, M. S., Mehta, S. & Zhang, J. Next-Generation Genetically Encoded Fluorescent Biosensors Illuminate Cell Signaling and Metabolism. Annual Review of Biophysics 53, 275–297 (2024). 10.1146/annurev-biophys-030722-021359

14 Alex, A. et al. Electroporated recombinant proteins as tools for in vivo functional complementation, imaging and chemical biology. eLife 8, e48287 (2019). 10.7554/eLife.48287

15 Nalbant, P., Hodgson, L., Kraynov, V., Toutchkine, A. & Hahn, K. M. Activation of endogenous Cdc42 visualized in living cells. Science 305, 1615–1619 (2004). 10.1126/science.1100367

16 Farrants, H. et al. A modular chemigenetic calcium indicator for multiplexed in vivo functional imaging. Nature Methods 21, 1916–1925 (2024). 10.1038/s41592-024-02411-6

17 Huppertz, M.-C. et al. Recording physiological history of cells with chemical labeling. Science 383, 890–897 (2024). doi:10.1126/science.adg0812

18 Toutchkine, A., Kraynov, V. & Hahn, K. Solvent-sensitive dyes to report protein conformational changes in living cells. Journal of the American Chemical Society 125, 4132–4145 (2003). 10.1021/ja0290882

19 MacNevin, C. J. et al. Environment-sensing merocyanine dyes for live cell imaging applications. Bioconjugate Chemistry 24, 215–223 (2013). 10.1021/bc3005073

20 Molla, A., Katz, S. & Demaille, J. G. in Current Topics in Membranes and Transport Vol. 25 (ed Felix Bronner) 147–180 (Academic Press, 1985).

21 Cohen, S. M. et al. Calmodulin shuttling mediates cytonuclear signaling to trigger experience-dependent transcription and memory. Nature Communications 9, 2451 (2018). 10.1038/s41467-018-04705-8

22 Zhang, M., Tanaka, T. & Ikura, M. Calcium-Induced Conformational Transition Revealed by the Solution Structure of Apo Calmodulin. Nat Struct Biol 2, 758–767 (1995). 10.1038/Nsb0995-758

23 Clow, A., Jenner, P., Theodorou, A. & Marsden, C. D. Striatal dopamine receptors become supersensitive while rats are given trifluoperazine for six months. Nature 278, 59–61 (1979). 10.1038/278059a0

24 Clow, A., Theodorou, A., Jenner, P. & Marsden, C. D. A comparison of striatal and mesolimbic dopamine function in the rat during 6-month trifluoperazine administration. Psychopharmacology 69, 227–233 (1980). 10.1007/BF00433087

25 Ujiié, Y. et al. Unique evolution of foraminiferal calcification to survive global changes. Science Advances 9, eadd3584 (2023). doi:10.1126/sciadv.add3584

26 Guamán-Guevara, F., Austin, H., Hicks, N., Streeter, R. & Austin, W. E. N. Impacts of ocean acidification on intertidal benthic foraminiferal growth and calcification. PLOS ONE 14, e0220046 (2019). 10.1371/journal.pone.0220046

27 Hallock, P., Lidz, B. H., Cockey-Burkhard, E. M. & Donnelly, K. B. Foraminifera as Bioindicators in Coral Reef Assessment and Monitoring: The FORAM Index. Environmental Monitoring and Assessment 81, 221–238 (2003). 10.1023/A:1021337310386

28 Mehl, B. P. et al. Live-cell biosensors based on the fluorescence lifetime of environment-sensing dyes. Cell Rep Methods 4, 100734 (2024). 10.1016/j.crmeth.2024.100734

29 Sathya, V. et al. Development of Optical Biosensor for the Detection of Glutamine in Human Biofluids Using Merocyanine Dye. J Fluoresc 32, 1389–1396 (2022). 10.1007/s10895-022-02937-y

30 Zhang, X. et al. ROS/RNS and Base Dual Activatable Merocyanine-Based NIR-II Fluorescent Molecular Probe for in vivo Biosensing. Angew Chem Int Ed Engl 60, 26337–26341 (2021). 10.1002/anie.202109728

31 Morley, J. O., Morley, R. M., Docherty, R. & Charlton, M. H. Fundamental Studies on Brooker’s Merocyanine. Journal of the American Chemical Society 119, 10192–10202 (1997). 10.1021/ja971477m

32 Han, W.-G. et al. A Theoretical Study of the UV/Visible Absorption and Emission Solvatochromic Properties of Solvent-Sensitive Dyes. ChemPhysChem 4, 1084–1094 (2003). 10.1002/cphc.200300801

33 Liu, T. et al. Density Functional Vertical Self-Consistent Reaction Field Theory for Solvatochromism Studies of Solvent-Sensitive Dyes. The Journal of Physical Chemistry A 108, 3545–3555 (2004). 10.1021/jp031062p

34 Toutchkine, A. et al. Experimental and DFT studies: novel structural modifications greatly enhance the solvent sensitivity of live cell imaging dyes. J Phys Chem A 111, 10849–10860 (2007). 10.1021/jp073197r

35 Toutchkine, A., Nguyen, D. V. & Hahn, K. M. Merocyanine dyes with improved photostability. Org Lett 9, 2775–2777 (2007). 10.1021/ol070780h

36 Massom, L., Lee, H. & Jarrett, H. W. Trifluoperazine Binding to Porcine Brain Calmodulin and Skeletal-Muscle Troponin-C. Biochemistry 29, 671–681 (1990). 10.1021/Bi00455a012

37 Vandonselaar, M., Hickie, R. A., Quail, W. & Delbaere, L. T. J. Trifluoperazine-induced conformational change in Ca2+-calmodulin. Nat Struct Biol 1, 795–801 (1994). 10.1038/nsb1194-795

38 Cook, W. J., Walter, L. J. & Walter, M. R. Drug binding by calmodulin: crystal structure of a calmodulin-trifluoperazine complex. Biochemistry 33, 15259–15265 (1994). 10.1021/bi00255a006

39 Vertessy, B. G. et al. Simultaneous Binding of Drugs with Different Chemical Structures to Ca2+-Calmodulin: Crystallographic and Spectroscopic Studies. Biochemistry 37, 15300–15310 (1998). 10.1021/bi980795a

40 Levin, R. M. & Weiss, B. Specificity of the binding of trifluoperazine to the calcium-dependent activator of phosphodiesterase and to a series of other calcium-binding proteins. Biochim Biophys Acta 540, 197–204 (1978). 10.1016/0304-4165(78)90132-0

41 Tsien, R. Y. A non-disruptive technique for loading calcium buffers and indicators into cells. Nature 290, 527–528 (1981). 10.1038/290527a0

42 Lavis, L. D. Ester bonds in prodrugs. ACS Chem Biol 3, 203–206 (2008). 10.1021/cb800065s

43 Rotman, B. & Papermaster, B. W. Membrane properties of living mammalian cells as studied by enzymatic hydrolysis of fluorogenic esters. Proc Natl Acad Sci U S A 55, 134–141 (1966). 10.1073/pnas.55.1.134

44 Schultz, C. Prodrugs of biologically active phosphate esters. Bioorg Med Chem 11, 885–898 (2003). 10.1016/s0968-0896(02)00552-7

45 Rosch, U., Yao, S., Wortmann, R. & Wurthner, F. Fluorescent H-aggregates of merocyanine dyes. Angew Chem Int Ed Engl 45, 7026–7030 (2006). 10.1002/anie.200602286

46 Wurthner, F., Yao, S., Debaerdemaeker, T. & Wortmann, R. Dimerization of merocyanine dyes. Structural and energetic characterization of dipolar dye aggregates and implications for nonlinear optical materials. Journal of the American Chemical Society 124, 9431–9447 (2002). 10.1021/ja020168f

47 Agafonov, M., Volkova, T., Kumeev, R., Chibunova, E. & Terekhova, I. Impact of pluronic F127 on aqueous solubility and membrane permeability of antirheumatic compounds of different structure and polarity. Journal of Molecular Liquids 274, 770–777 (2019). 10.1016/j.molliq.2018.11.060

48 Hahn, K., DeBiasio, R. & Taylor, D. L. Patterns of elevated free calcium and calmodulin activation in living cells. Nature 359, 736–738 (1992). 10.1038/359736a0

49 McNeil, P. L., McKenna, M. P. & Taylor, D. L. A transient rise in cytosolic calcium follows stimulation of quiescent cells with growth factors and is inhibitable with phorbol myristate acetate. J Cell Biol 101, 372–379 (1985). 10.1083/jcb.101.2.372

50 Byron, K. L. & Villereal, M. L. Mitogen-induced [Ca2+]i changes in individual human fibroblasts: Image analysis reveals asynchronous responses which are characteristic for different mitogens*. Journal of Biological Chemistry 264, 18234–18239 (1989). 10.1016/S0021-9258(19)84702-6

51 Tucker, R. W. & Fay, F. S. Distribution of intracellular free calcium in quiescent BALB/c 3T3 cells stimulated by platelet-derived growth factor. Eur J Cell Biol 51, 120–127 (1990).

52 Li, C. J. et al. Dynamic redistribution of calmodulin in HeLa cells during cell division as revealed by a GFP-calmodulin fusion protein technique. J Cell Sci 112 (Pt 10), 1567–1577 (1999). 10.1242/jcs.112.10.1567

53 Nakai, J., Ohkura, M. & Imoto, K. A high signal-to-noise Ca2+ probe composed of a single green fluorescent protein. Nature Biotechnology 19, 137–141 (2001). 10.1038/84397

54 Klymchenko, A. S. Solvatochromic and Fluorogenic Dyes as Environment-Sensitive Probes: Design and Biological Applications. Accounts of Chemical Research 50, 366–375 (2017). 10.1021/acs.accounts.6b00517

55 Datta, R., Heaster, T. M., Sharick, J. T., Gillette, A. A. & Skala, M. C. Fluorescence lifetime imaging microscopy: fundamentals and advances in instrumentation, analysis, and applications. J Biomed Opt 25, 1–43 (2020). 10.1117/1.Jbo.25.7.071203

56 Vallmitjana, A., Torrado, B. & Gratton, E. Phasor-based image segmentation: machine learning clustering techniques. Biomed Opt Express 12, 3410–3422 (2021). 10.1364/boe.422766

57 Vallmitjana, A. et al. GSLab: open-source platform for advanced phasor analysis in fluorescence microscopy. Bioinformatics 41 (2025). 10.1093/bioinformatics/btaf162

58 Murthy, K. S. Signaling for contraction and relaxation in smooth muscle of the gut. Annual review of physiology 68, 345–374 (2006). 10.1146/annurev.physiol.68.040504.094707

59 Bowser, S. S., McGee-Russell, S. M. & Rieder, C. L. Digestion of prey in foraminifera is not anomalous: A correlation of light microscopic, cytochemical, and hvem technics to study phagotrophy in two allogromiids. Tissue and Cell 17, 823–839 (1985). 10.1016/0040-8166(85)90039-4

60 Bowser, S. S., Travis, J. L. & Rieder, C. L. Microtubules associate with actin-containing filaments at discrete sites along the ventral surface of Allogromia reticulopods. J Cell Sci 89 (Pt 3), 297–307 (1988). 10.1242/jcs.89.3.297

61 Ohno, Y., Fujita, K., Toyofuku, T. & Nakamaura, T. Cytological Observations of the Large Symbiotic Foraminifer Amphisorus kudakajimensis Using Calcein Acetoxymethyl Ester. PLOS ONE 11, e0165844 (2016). 10.1371/journal.pone.0165844

62 Travis, J. L., Kenealy, J. F. & Allen, R. D. Studies on the motility of the foraminifera. II. The dynamic microtubular cytoskeleton of the reticulopodial network of Allogromia laticollaris. Journal of Cell Biology 97, 1668–1676 (1983). 10.1083/jcb.97.6.1668

63 Tyszka, J. et al. Form and function of F-actin during biomineralization revealed from live experiments on foraminifera. Proceedings of the National Academy of Sciences 116, 4111–4116 (2019). doi:10.1073/pnas.1810394116

64 Miller, D. J. & Smith, G. L. EGTA purity and the buffering of calcium ions in physiological solutions. American Journal of Physiology-Cell Physiology 246, C160–C166 (1984). 10.1152/ajpcell.1984.246.1.C160

65 Liu, B. et al. Biosensors based on peptide exposure show single molecule conformations in live cells. Cell 184, 5670–5685.e5623 (2021). 10.1016/j.cell.2021.09.026

66 Abramson, J. et al. Accurate structure prediction of biomolecular interactions with AlphaFold 3. Nature 630, 493–500 (2024). 10.1038/s41586-024-07487-w

67 Forli, S. et al. Computational protein–ligand docking and virtual drug screening with the AutoDock suite. Nature Protocols 11, 905–919 (2016). 10.1038/nprot.2016.051

68 Erden, M., Devkota, K., Varghese, L., Cowen, L. & Singh, R. Learning a CoNCISE language for small-molecule binding. bioRxiv, 2025.2001.2008.632039 (2025). 10.1101/2025.01.08.632039

69 Singh, R., Sledzieski, S., Bryson, B., Cowen, L. & Berger, B. Contrastive learning in protein language space predicts interactions between drugs and protein targets. Proceedings of the National Academy of Sciences 120, e2220778120 (2023). doi:10.1073/pnas.2220778120

70 Yang, W. et al. The past, present and future of de novo protein design. Nature 652, 1139–1152 (2026). 10.1038/s41586-026-10328-7

71 Hart, K. M. et al. Designing small molecules to target cryptic pockets yields both positive and negative allosteric modulators. PLOS ONE 12, e0178678 (2017). 10.1371/journal.pone.0178678

72 Redij, T., Chaudhari, R., Li, Z., Hua, X. & Li, Z. Structural Modeling and in Silico Screening of Potential Small-Molecule Allosteric Agonists of a Glucagon-like Peptide 1 Receptor. ACS Omega 4, 961–970 (2019). 10.1021/acsomega.8b03052

73 Hahn, K. & Toutchkine, A. Live-cell fluorescent biosensors for activated signaling proteins. Current Opinion in Cell Biology 14, 167–172 (2002). 10.1016/S0955-0674(02)00313-7

74 Liu, S., Su, Y., Lin, M. Z. & Ronald, J. A. Brightening up Biology: Advances in Luciferase Systems for in Vivo Imaging. ACS Chemical Biology 16, 2707–2718 (2021). 10.1021/acschembio.1c00549

75 Zhang, J., Ma, Y., Taylor, S. S. & Tsien, R. Y. Genetically encoded reporters of protein kinase A activity reveal impact of substrate tethering. Proceedings of the National Academy of Sciences 98, 14997–15002 (2001). doi:10.1073/pnas.211566798

76 Ting, A. Y., Kain, K. H., Klemke, R. L. & Tsien, R. Y. Genetically encoded fluorescent reporters of protein tyrosine kinase activities in living cells. Proceedings of the National Academy of Sciences 98, 15003–15008 (2001). doi:10.1073/pnas.211564598

77 Swanson, J. in Methods in Cell Biology Vol. 29 (eds Yu-Li Wang, D. Lansing Taylor, & K. W. Jeon) 137–151 (Academic Press, 1988).

78 Zuidscherwoude, M. et al. Calmodulin regulates TRPV5 intracellular trafficking and plasma membrane abundance. The Journal of Physiology 602, 6871–6888 (2024). 10.1113/JP286182

79 Villalobo, A., Ruano, M. J., Palomo-Jiménez, P. I., Li, H. & Martín-Nieto, J. in Calcium: The Molecular Basis of Calcium Action in Biology and Medicine (eds Roland Pochet et al.) 287–303 (Springer Netherlands, 2000).

80 Walsh, M. P. Calmodulin and the regulation of smooth muscle contraction. Mol Cell Biochem 135, 21–41 (1994). 10.1007/bf00925958

81 Schoenmakers, T. J., Visser, G. J., Flik, G. & Theuvenet, A. P. CHELATOR: an improved method for computing metal ion concentrations in physiological solutions. Biotechniques 12, 870–874, 876–879 (1992).

82 Edelstein, A. D. et al. Advanced methods of microscope control using μManager software. J Biol Methods 1 (2014). 10.14440/jbm.2014.36

83 Schindelin, J. et al. Fiji: an open-source platform for biological-image analysis. Nature Methods 9, 676–682 (2012). 10.1038/nmeth.2019

84 Kuboniwa, H. et al. Solution structure of calcium-free calmodulin. Nat Struct Biol 2, 768–776 (1995). 10.1038/nsb0995-768

85 Chattopadhyaya, R., Meador, W. E., Means, A. R. & Quiocho, F. A. Calmodulin structure refined at 1.7 Å resolution. Journal of Molecular Biology 228, 1177–1192 (1992). 10.1016/0022-2836(92)90324-D

