## Supplementary Figures and Movie Legends for "A membrane-permeable small molecule biosensor accesses intractable cells and animals without genetic manipulation"

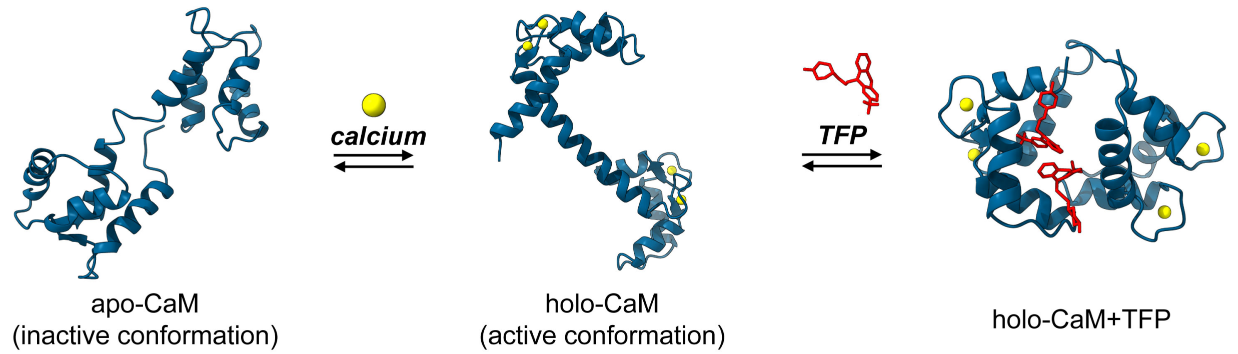


**Supplementary Figure 1. Inactive versus active calmodulin conformation.** Upon binding $Ca^{2+}$, calmodulin (PDB: 1CFD, apo calmodulin)[1] undergoes a shift from a less ordered conformation to a dumbbell shaped holo conformation (PDB: 1CLL)[2]. Holo-calmodulin binds to the small molecule trifluoperazine (PDB: 1A29)[3]. Up to four TFP binding sites of widely differing affinity have been identified on calmodulin[4]. Two TFP molecules bind to the hydrophobic pockets in the heads of the “dumbbell” at 1-2 μM affinity. These may induce a conformational change that can allow two additional TFP molecules to bind, but these binding interactions are less well defined.

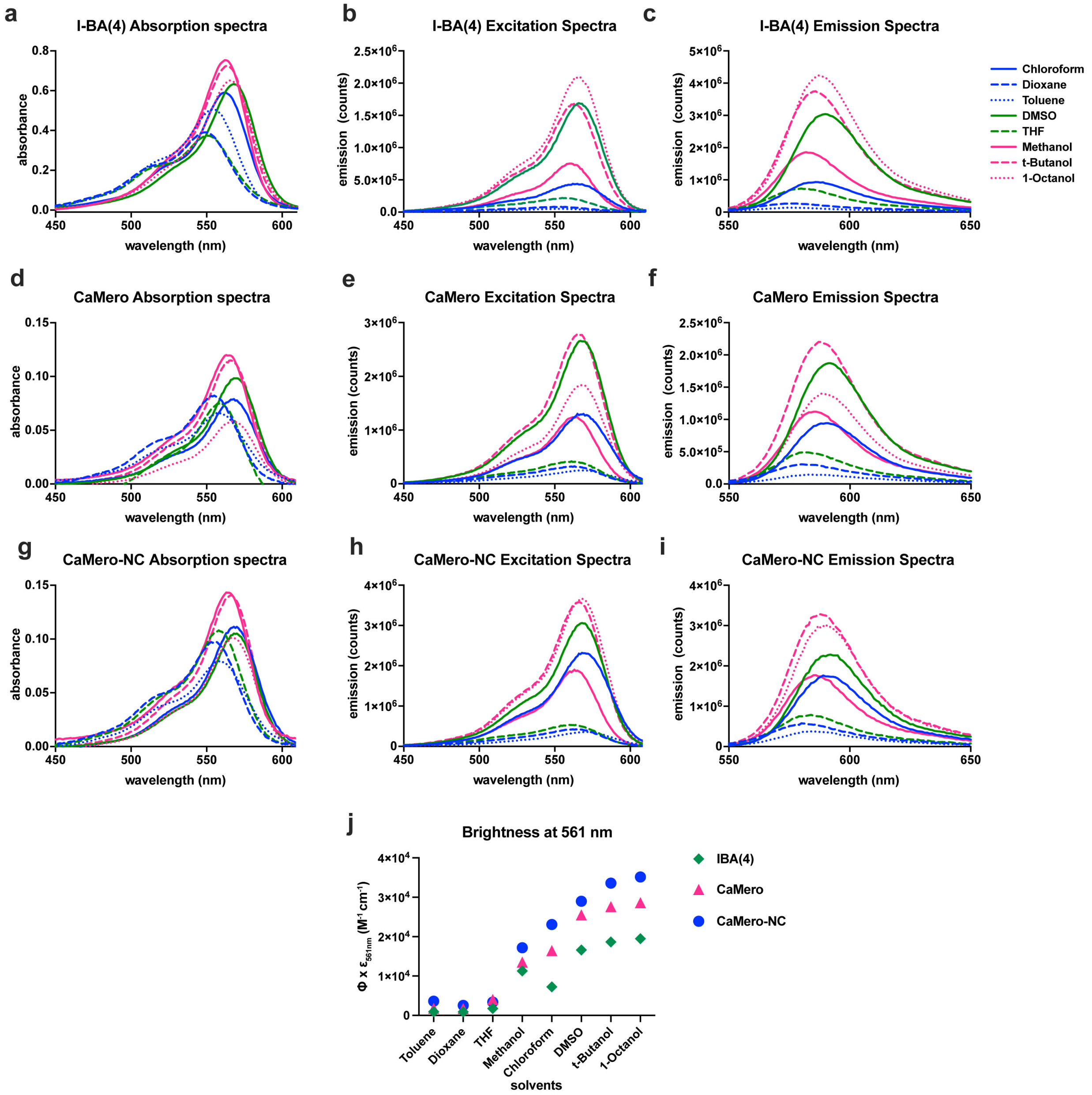


**Supplementary Figure 2. Photophysical properties of I-BA(4), CaMero and CaMero-NC in organic solvents.** (**a–c**) Absorption, excitation and emission spectra of I-BA(4). (**d–f**) Absorption, excitation and emission spectra of CaMero. (**g–i**) Absorption, excitation and emission spectra of CaMero-NC. (**j**) Brightness (extinction coefficient at 561 nm x quantum yield) of I-BA(4), CaMero, and CaMero-NC in different solvents.

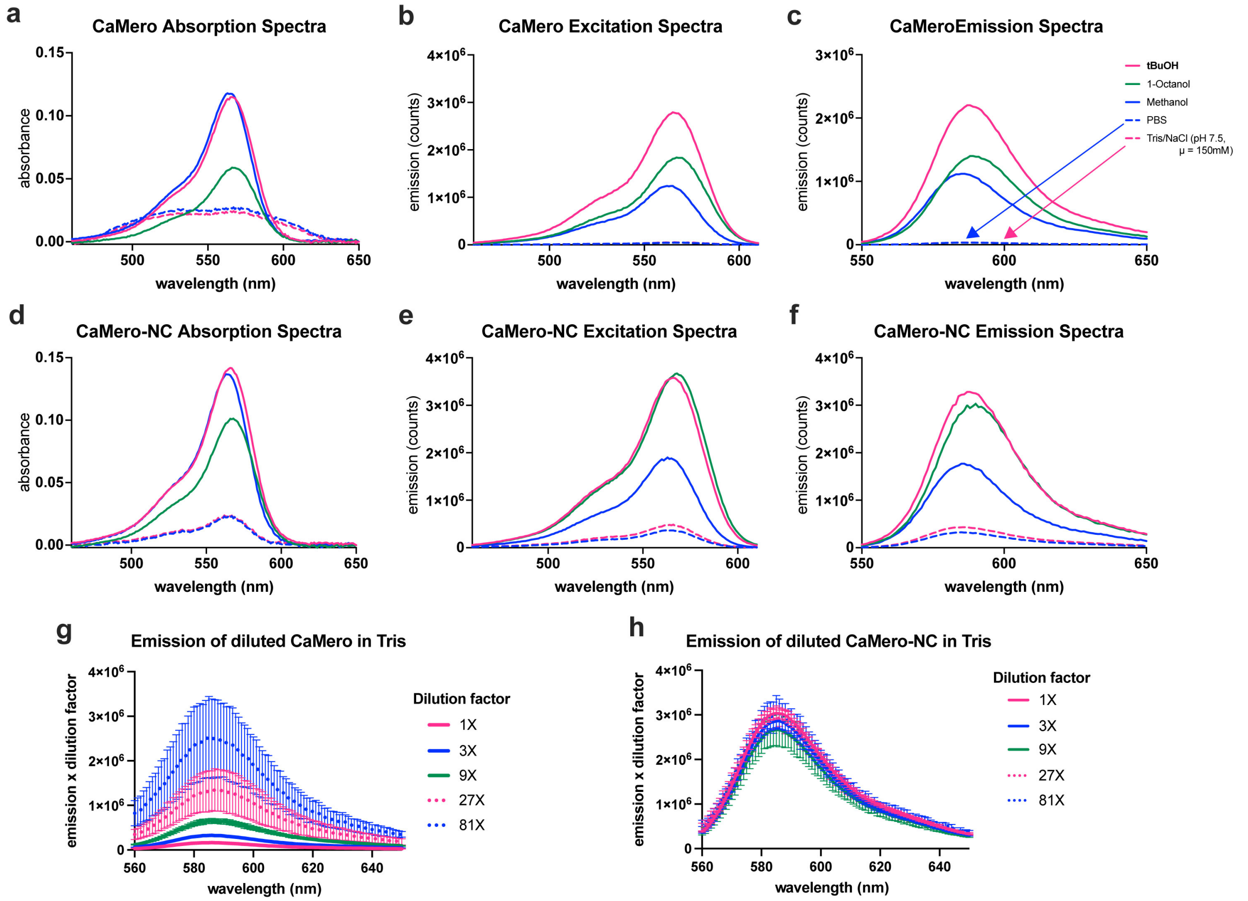


**Supplementary Figure 3. Photophysical properties of CaMero and CaMero-NC in aqueous solvents and alcohols.** (**a–c**) Absorption, excitation, and emission spectra of CaMero. (**d–f**) Absorption, excitation, and emission spectra of CaMero-NC. (**g**) Emission spectra of serially diluted samples of CaMero in Tris buffer; emission counts are multiplied by the corresponding dilution factor. (**h**) Emission spectra of serially diluted samples of CaMero-NC in Tris buffer (1X = 2 μM); emission counts are multiplied by the corresponding dilution factor.

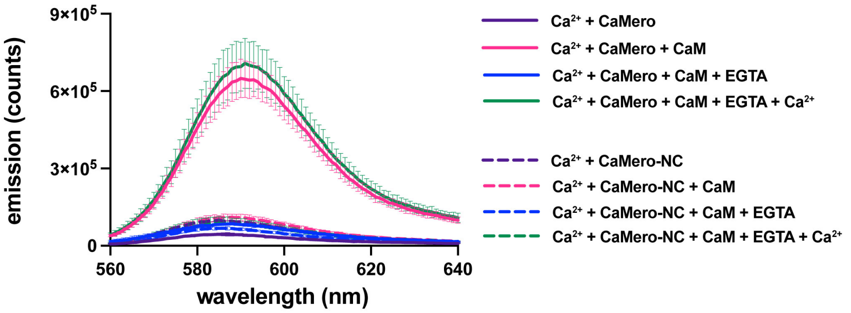


**Supplementary Figure 4. Reversibility and $Ca^{2+}$-dependence of CaMero and CaMero-NC binding to calmodulin.** (**a**) Emission spectra of CaMero and CaMero-NC recorded in the absence and presence of active/inactive calmodulin.

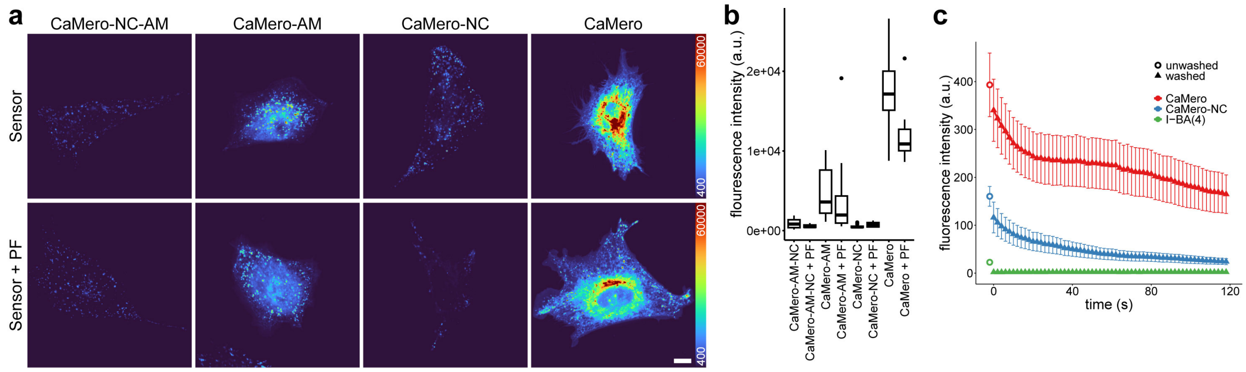


**Supplementary Figure 5. Comparison of probes and determination of optimal imaging conditions.** (**a**) Use of Pluronic F-127 (PF) or addition of acetoxymethyl (AM) to the dyes did not increase cell loading. Pluronic F-127 was added to the initial DMSO stock solutions of dyes to give a final concentration in the incubation solutions of < 0.01%. (**b**) Fluorescence intensity per unit area for at least 15 cells per condition. (**c**) Mean fluorescence intensity before (circles) and after (triangles) washing out the biosensors and I-BA(4) dye, corrected for background.

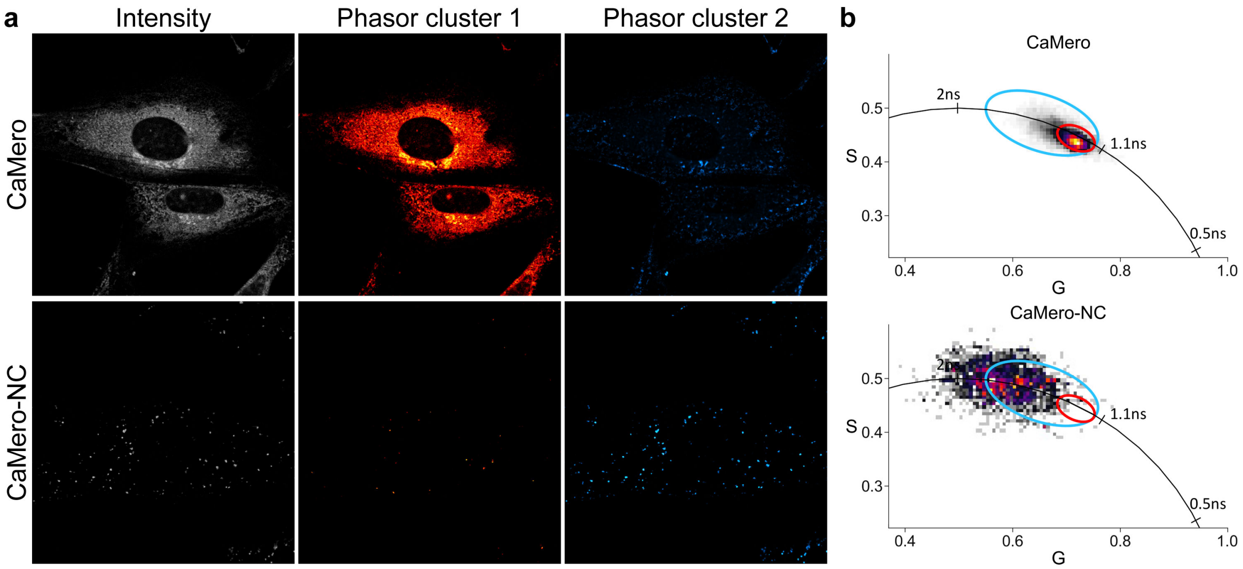


**Supplementary Figure 6. Fluorescence lifetime of CaMero changes upon binding calmodulin.** (**a**) FLIM imaging of CaMero and CaMero-NC. (**b**) Phasor analysis of CaMero shows a distinct lifetime when bound to calmodulin while CaMero-NC is much more diffuse with a longer lifetime.

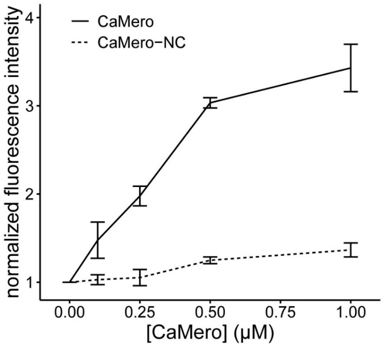


**Supplementary Figure 7. Determination of optimal biosensor concentration.** Intracellular intensity of CaMero increased with extracellular concentration of CaMero in the medium during incubation. CaMero-NC showed minimal change.

**Supplementary Video 1. Calmodulin activity reported by CaMero in MEF cells responding to serum stimulation.** MEF cells incubated with CaMero or CaMero-NC probe were serum starved and then treated with FBS. Images were scaled identically, with scale selected to prevent camera saturation at peak CaMero intensity.

**Supplementary Video 2. Calmodulin activity in a foraminifera treated with EGTA-AM and seawater.** A foraminifera moving along the coverslip after recovery from EGTA-AM treatment by adding additional seawater. Brightfield (grey) is shown on the left while CaMero (cyan) and chlorophyll (magenta) fluorescence are shown on the right.

### REFERENCES


1 Kuboniwa, H. *et al.* Solution structure of calcium-free calmodulin. *Nat Struct Biol* **2**, 768–776 (1995). https://doi.org/10.1038/nsb0995-768

2 Chattopadhyaya, R., Meador, W. E., Means, A. R. & Quiocho, F. A. Calmodulin structure refined at 1.7 Å resolution. *Journal of Molecular Biology* **228**, 1177–1192 (1992). https://doi.org/10.1016/0022-2836(92)90324-D

3 Vertessy, B. G. *et al.* Simultaneous Binding of Drugs with Different Chemical Structures to Ca2+-Calmodulin: Crystallographic and Spectroscopic Studies. *Biochemistry* **37**, 15300–15310 (1998). https://doi.org/10.1021/bi980795a

4 Vandonselaar, M., Hickie, R. A., Quail, W. & Delbaere, L. T. J. Trifluoperazine-induced conformational change in Ca2+-calmodulin. *Nat Struct Biol* **1**, 795–801 (1994). https://doi.org/10.1038/nsb1194-795
