## Supplementary Methods for "A membrane-permeable small molecule biosensor accesses intractable cells and animals without genetic manipulation"

### GENERAL METHODS AND MATERIALS

All commercially available materials were used as received. Reaction progress was monitored via thin-layer chromatography (TLC) on pre-coated aluminum-backed plates (silica gel 60 Å $F_{254}$). Flash chromatography was carried out with silica gel 60 Å (230 - 400 mesh) and automated chromatography was performed on a Teledyne-Isco Combiflash Rf system or BUCHI Pure System. Unless otherwise stated, organic extracts were dried over commercially available magnesium sulfate, filtered and the filtrate was concentrated via rotary evaporation. Mass spectra were obtained on a Hewlett-Packard 1100 high-performance liquid chromatograph equipped with an 1100 mass-selective detector (MS-ESI) or a Thermo Scientific Q Exactive HF-X mass spectrometer with direct infusion. UV-visible spectra were obtained with a Hewlett-Packard 8453 diode array spectrophotometer or Agilent Cary 60 UV-Vis spectrophotometer. Emission and excitation spectra were obtained using a Spex Fluorolog 2 spectrofluorometer at 23 °C. High performance liquid chromatography was performed on a Shimadzu Prominence system using a Phenomenex C18 preparative column (250 x 21.2 mm, 15 μM particle size) and elution at 8 mL/min with a gradient of 10% solvent B ($H_2O$/acetonitrile 5:95, TFA 0.05%) 90% solvent A ($H_2O$/acetonitrile 95:5, 0.05% TFA) for 2 min, increasing to 90% solvent B over 30 min and held for a total of 45 min. $^1H$ and $^{13}C$ NMR spectra were recorded on a Varian Inova 400 MHz or Bruker-500 spectrometer using deuterated solvents and referenced to the residual solvent peak (for $CDCl_3$ = $^1H$ δ 7.26 ppm, $^{13}C$ δ 77.16 ppm; for DMSO-$d_6$ = $^1H$ δ 2.50 ppm, $^{13}C$ δ 39.52 ppm). All operations with dyes were performed under dim light. **Mero76** was synthesized as previously described[1]. **CaMero-AM** and **CaMero-AM-NC** were synthesized as described in **Scheme S1**. **I-BA(4)-NHS** was synthesized as described in **Scheme S2**. **CaMero** and **CaMero-NC** were synthesized as described in **Scheme S3**.

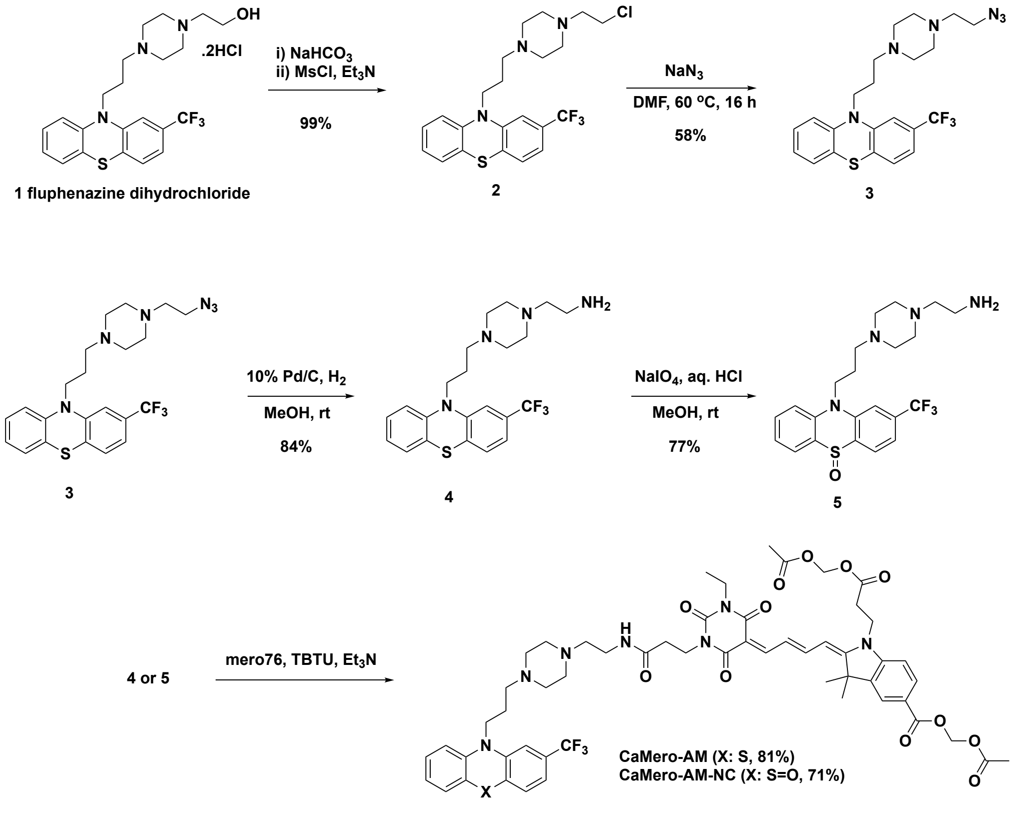


**Scheme S1**. Synthesis of **CaMero-AM** and **CaMero-AM-NC**.

**10-(3-(4-(2-Chloroethyl)piperazin-1-yl)propyl)-2-(trifluoromethyl)-10H-phenothiazine (2)**. Fluphenazine dihydrochloride (**1**, 1.00 g, 1.96 mmol) was diluted in saturated aqueous sodium bicarbonate (50 mL) and stirred for 10 min. The aqueous layer was extracted with $CH_2Cl_2$ (3 x 50 mL). Organic layers were combined, washed with saturated aq. NaCl, dried with $MgSO_4$, and filtered. The filtrate was concentrated via rotary evaporation to give 0.85 g pale amber oil (quant.) that was carried forward directly to the next step.

The crude product (0.845 g, 1.93 mmol) was diluted in $CH_2Cl_2$ (25 mL) in an oven dried 100 mL round-bottom flask. The flask was sealed with a rubber septum, flushed with argon, and chilled in an ice bath. Triethylamine (0.417 mL, 2.99 mmol) was added, followed by dropwise addition of methanesulfonyl chloride (0.233 mL, 2.99 mmol). The mixture was stirred for 1 h at 0 °C, allowed to equilibrate to 25 ºC and stirred for 1.5 h. Water was added (25 mL) and the organic layer was separated, dried with $MgSO_4$ and filtered. The filtrate was concentrated via rotary evaporation to yield 0.893 g (quant.) pale amber oil. The product was carried forward without further purification. $^1H$ NMR (400 MHz, $CDCl_3$) δ 7.24 – 7.08 (m, 4H), 7.03 (s, 1H), 6.97 – 6.88 (m, 2H), 3.95 (t, *J* = 6.8 Hz, 2H), 3.56 (t, *J* = 7.1 Hz, 2H), 2.70 (t, *J* = 7.1 Hz, 2H), 2.62 – 2.25 (m, 10H), 1.98 – 1.87 (m, 2H). $^{13}C$ NMR (100 MHz, $CDCl_3$) δ 145.9, 144.5, 130.1, 129.7 (q, *J* = 32 Hz), 127.8, 127.7, 127.6, 124.3 (q, *J* = 270 Hz), 124.2, 123.2, 119.1 (q, *J* = 3.9 Hz), 116.1, 112.1 (q, *J* = 3.8 Hz), 59.9, 55.5, 53.3 (four C), 45.4, 41.0, 24.3. MS-ESI *m/z* 456.2 ($[M + H]^+$ requires 456.1).

**10-(3-(4-(2-Azidoethyl)piperazin-1-yl)propyl)-2-(trifluoromethyl)-10H-phenothiazine (3)**. Compound **2** (1.52 g, 3.33 mmol) was added to a 50 mL oven dried round-bottom flask and diluted in DMF (20.0 mL). Sodium iodide (0.599 g, 3.99 mmol) was added, followed by sodium azide (0.433 g, 6.67 mmol). The flask was sealed with a septum, argon flushed, and the reaction mixture was heated to 60 °C for 16 h. Aqueous LiCl (15% w/v, 50 mL) was added and the aqueous layer was extracted with EtOAc (3 x 50 mL). The organic layers were combined, washed with water (200 mL) and brine (200 mL), dried with $Na_2SO_4$, filtered, and concentrated. The crude product was suspended in a minimum amount of $CH_2Cl_2$ and loaded onto a 24 g $SiO_2$ column. The column was eluted with 0 – 3% MeOH in $CH_2Cl_2$ over 20 min. Main product containing fractions were combined, concentrated, and dried under vacuum to give 0.890 g (58%) pale amber oil. $^1H$ NMR (400 MHz, $CDCl_3$) δ 7.22 – 7.09 (m, 4H), 7.04 (s, 1H), 6.97 – 6.88 (m, 2H), 3.96 (t, J = 6.8 Hz, 1H), 3.32 (t, J = 6.1 Hz, 1H), 2.56 (t, J = 6.1 Hz, 2H), 2.54 – 2.20 (m, 8H), 1.97 – 1.88 (m, 2H). $^{13}C$ NMR (100 MHz, $CDCl_3$) δ 145.9, 144.5, 130.0, 129.7 (q, *J* = 32 Hz), 127.8, 127.7, 127.6, 124.3 (q, *J* = 271 Hz), 124.2, 123.2, 119.1 (q, *J* = 3.9 Hz), 116.1, 112.1 (q, *J* = 3.8 Hz), 57.2, 55.5, 53.3 (four C), 48.4, 45.5, 24.3. MS-ESI *m/z* 463.2 ($[M + H]^+$ requires 463.2).

**2-(4-(3-(2-(Trifluoromethyl)-10H-phenothiazin-10-yl)propyl)piperazin-1-yl)ethan-1-amine (4)**. Palladium on carbon (10 wt. %, 0.060 g) was added to a 50 mL flame dried two-neck round-bottom flask. One neck of the flask was sealed with a rubber septum fitted with an argon line and the other with a glass stopper. The flask was evacuated and argon flushed two times. Compound **3** (0.305 g, 0.659 mmol) was diluted in anhydrous methanol (10 mL) and added to the reaction flask. The glass stopper was replaced with a hydrogen containing balloon. The argon line was removed and the flask was flushed with hydrogen (2 times). The balloon was left open to the system and the reaction mixture was stirred at 25 ºC for 16 h. The reaction mixture was filtered through a sand-topped Celite column and the column was rinsed with methanol (75 mL). The filtrate was concentrated, diluted in $CH_2Cl_2$, dried with $Na_2SO_4$, and filtered with $CH_2Cl_2$ washes. The filtrates were concentrated via rotary evaporation and loaded in a minimum amount of $CH_2Cl_2$ onto a 12 g $SiO_2$ column and eluted with 0 – 10% MeOH (containing 1% $NH_3$) in $CH_2Cl_2$ over 25 min. Main peak containing fractions were combined, concentrated, and dried under vacuum to give 0.242 g (84%) colorless oil. $^{1}$H NMR (400 MHz, $CDCl_3$) δ 7.22 – 7.07 (m, 4H), 7.03 (s, 1H), 6.97 – 6.88 (m, 2H), 3.95 (t, *J* = 6.8 Hz, 2H), 2.76 (t, *J* = 6.2 Hz, 2H), 2.63 – 2.20 (m, 12H), 2.01 – 1.85 (m, 2H), 1.41 (bs, 2H). $^{13}$C NMR (100 MHz, $CDCl_3$) δ 145.9, 144.5, 130.0, 129.7 (q, J = 32.1 Hz), 127.8, 127.7, 127.6, 124.3 (q, J = 271 Hz), 124.2, 123.2, 119.1 (q, J = 3.9 Hz), 116.1, 112.0 (q, J = 3.8 Hz), 61.2, 55.6, 53.5 (two C), 53.4 (two C), 45.5, 39.0, 24.3. MS-ESI *m/z* 437.2 ($[M + H]^+$ requires 437.2).

**10-(3-(4-(2-Aminoethyl)piperazin-1-yl)propyl)-2-(trifluoromethyl)-10H-phenothiazine 5-oxide (5)**. Compound **4** (0.050 g, 0.11 mmol) was diluted in MeOH (1.0 mL) in a 10 mL round-bottom flask open to air. Sodium periodate (0.028 g, 0.132 mmol) was dissolved in a 3:1 MeOH/1.0 M aq. HCl solution (2.0 mL) with sonication and added to the reaction flask quickly dropwise. The reaction mixture was stirred for 4 h at 25 ºC. Saturated aq. $NaHCO_3$ (2.0 mL) was added and the mixture was concentrated. Water was added (10 mL) and the aqueous layer was extracted with $CH_2Cl_2$ (10 mL x 4). Organic layers were combined, washed with saturated aq. NaCl, dried with $MgSO_4$, filtered, and concentrated via rotary evaporation to give 0.040 g (77%) clear oil that foamed on drying. The product required no further purification. $^{1}$H NMR (400 MHz, $CDCl_3$) δ 8.04 (d, *J* = 8.0 Hz, 1H), 7.95 (dd, *J* = 7.7, 1.5 Hz, 1H), 7.72 (s, 1H), 7.68 – 7.62 (m, 1H), 7.59 (d, *J* = 8.3 Hz, 1H), 7.47 (d, *J* = 8.0 Hz, 1H), 7.30 (t, *J* = 7.4 Hz, 1H), 4.50 – 4.31 (m, 2H), 2.79 (t, *J* = 6.2 Hz, 2H), 2.69 – 2.27 (m, 12H), 2.12 – 1.99 (m, 2H), 1.63 (bs, 2H). $^{13}$C NMR (100 MHz, $CDCl_3$) δ 138.7, 138.1, 134.4 (q, *J* = 32.3 Hz), 133.1, 132.2, 131.5, 127.2, 124.6, 123.5 (q, *J* = 272 Hz), 122.6, 117.9 (q, *J* = 3.5 Hz), 116.3, 113.2 (q, *J* = 4.0 Hz), 61.1, 54.6, 53.4 (two C), 53.3 (two C), 45.6, 38.8, 23.9. MS-ESI *m/z* 453.2 ($[M + H]^+$ requires 453.2).

**CaMero-AM**. Compound **mero76** (10.0 mg, 0.015 mmol), compound **4** (9.6 mg, 0.022 mmol), and TBTU (7.0 mg, 0.022 mmol) were added to a 5 mL conical vial with spin vane and dissolved in DMF (0.70 mL), followed by addition of $Et_3N$ (3.1 µL, 0.022 mmol). The vial was capped and covered in foil and the reaction was stirred at 25 ºC for 16 h. The reaction mixture was submitted to preparative HPLC. Main product containing fractions were combined, concentrated, and lyophilized. Isolated 13.0 mg (81%) dark blue powdery solid. $^{1}$H NMR (400 MHz, DMSO) δ 8.24 (t, *J* = 12.2 Hz, 1H), 8.20 – 8.04 (m, 2H), 7.99 (s, 1H), 7.94 (d, *J* = 8.4 Hz, 1H), 7.81 (dd, *J* = 24.9, 12.9 Hz, 1H), 7.39 (d, *J* = 7.9 Hz, 1H), 7.32 – 7.26 (m, 4H), 7.22 (d, *J* = 7.5 Hz, 1H), 7.10 (dd, *J* = 7.9, 2.7 Hz, 1H), 7.03 (t, *J* = 7.4 Hz, 1H), 6.22 (dd, *J* = 15.6, 13.1 Hz, 1H), 5.93 (s, 2H), 5.62 (d, *J* = 2.6 Hz, 2H), 4.30 – 4.15 (m, 2H), 4.09 – 3.97 (m, 4H), 3.90 – 3.80 (m, 2H), 3.35 – 3.22 (m, 2H), 3.13 – 2.84 (m, 4H), 2.81 (t, *J* = 6.3 Hz, 2H), 2.37 (t, *J* = 6.4 Hz, 2H), 2.10 (s, 3H), 2.05 – 1.95 (m, 2H), 2.01 (d, *J* = 2.2 Hz, 3H), 1.63 (s, 6H), 1.10 (t, *J* = 7.0 Hz, 3H). HR-FTMS *m/z* 1102.4218 ($[M + H]^+$ requires 1102.4208).

**CaMero-AM-NC**. Compound **mero76** (10.2 mg, 0.015 mmol), compound **5** (10.2 mg, 0.023 mmol), and TBTU (7.2 mg, 0.022 mmol) were added to a 5 mL conical vial with spin vane and dissolved in DMF (0.70 mL), followed by addition of $Et_3N$ (3.2 µL, 0.023 mmol). The vial was capped and covered in foil and the reaction was stirred at 25 ºC for 16 h. The reaction mixture was submitted to preparative HPLC. Main product containing fractions were combined, concentrated, and lyophilized. Isolated 12.0 mg (71%) dark blue powdery solid. $^{1}$H NMR (400 MHz, DMSO) δ 8.28 – 7.90 (m, 6H), 7.98 (s, 1H), 7.97 – 7.89 (m, 1H), 7.86 – 7.73 (m, 3H), 7.61 (d, *J* = 8.1 Hz, 1H), 7.38 (t, *J* = 7.4 Hz, 1H), 7.29 (t, *J* = 9.0 Hz, 1H), 6.29 – 6.15 (m, 1H), 5.93 (s, 2H), 5.62 (d, *J* = 3.4 Hz, 2H), 4.65 – 4.60 (m, 2H), 4.30 – 4.17 (m, 2H), 4.02 (t, *J* = 6.5 Hz, 2H), 3.90 – 3.78 (m, 2H), 3.37 – 3.22 (m, 2H), 3.05 – 2.75 (m, 6H), 2.37 (t, *J* = 6.4 Hz, 2H), 2.10 (s, 3H), 2.09 – 2.02 (m, 2H), 2.00 (d, *J* = 2.8 Hz, 3H), 1.62 (s, 6H), 1.10 (t, *J* = 7.0 Hz, 3H). HR-FTMS *m/z* 1118.4162 ($[M + H]^+$ requires 1118.4157).

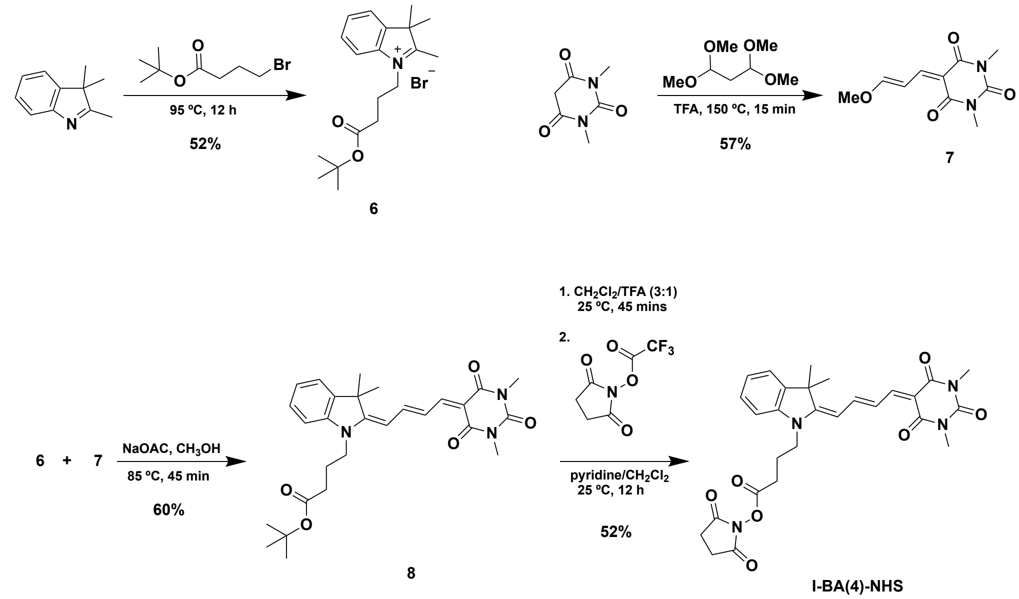


**Scheme S2**. Synthesis of **I-BA(4)-NHS**

**1-(4-(tert-Butoxy)-4-oxobutyl)-2,3,3-trimethyl-3H-indol-1-ium bromide (6).** 2,3,3-Trimethylindolenine (2.00 mL, 12.0 mmol), and tert-butyl 4-bromobutanoate (3.00 mL, 18.0 mmol) were mixed in a pressure tube. The resulting mixture was sonicated for a minute and then heated at 95 ºC for 12 h. The flask was cooled in an ice-bath and the contents were mixed with diethyl ether (20 mL). The resulting residue was separated and washed with diethyl ether to obtain the title compound as a pale pink solid (2.40 g, 52% yield). $^{1}$H NMR ($CDCl_3$, 500 MHz) δ ppm 7.95 (d, *J* = 7.5 Hz, 1H), 7.63-7.54 (m, 3H), 4.81 (t, *J* = 8.0 Hz, 2H), 3.17 (s, 3H), 2.61 (t, *J* = 6.0 Hz, 2H), 2.24-2.19 (m, 2H), 1.66 (s, 6H), 1.39 (s, 9H); $^{13}$C NMR ($CDCl_3$, 125 MHz) δ ppm 196.0, 171.8, 141.5, 141.3, 130.1, 129.7, 123.1, 115.8, 81.5, 54.6, 49.3, 31.7, 28.0, 23.2, 22.8, 17.0; HRMS (positive mode) obsd 302.2115, calcd 302.2115 [$M^+$, M = $C_{19}H_{28}NO_2$].

**(*E*)-5-(3-Methoxyallylidene)-1,3-dimethylpyrimidine-2,4,6(*1H,3H,5H*)-trione (7).** A mixture of 1,3-dimethylpyrimidine-2,4,6(1*H*,3*H,5H*)-trione (1.56 g, 10.0 mmol), and 1,1,3,3-tetramethoxypropane (10.0 mL) was treated with trifluoroacetic acid (100 µL) in a pressure tube. The resulting mixture was heated at 150 ºC for 15 min. The content was cooled in an ice bath and mixed with hexane (20 mL). The resulting residue was separated and washed with hexane to obtain the title compound as a dark orange solid (1.29 g, 57% yield). $^{1}$H NMR ($CDCl_3$, 500 MHz) δ ppm 8.10 (d, *J* = 12.0 Hz, 1H), 7.53 (d, *J* = 12.0 Hz, 1H), 7.48 (t, *J* = 12.0 Hz, 1H), 3.95 (s, 3H), 3.37 (s, 3H), 3.35 (s, 3H); $^{13}$C NMR ($CDCl_3$, 125 MHz) δ ppm 171.6, 162.6, 162.3, 157.7, 151.7, 110.3, 105.7, 58.7, 28.5, 27.8; HRMS (positive mode) obsd 225.0869, calcd 225.0870 [$M+H^+$, M = $C_{10}H_{12}N_2O_4$].

**tert-Butyl 4-((E)-2-((E)-4-(1,3-dimethyl-2,4,6-trioxotetrahydropyrimidin-5(2H)-ylidene)but-2-en-1-ylidene)-3,3-dimethylindolin-1-yl)butanoate (8).** Compound **6** (300 mg, 0.800 mmol), compound **7** (300 mg, 1.30 mmol) and sodium acetate (166 mg, 2.00 mmol) were dissolved in methanol (5.0 mL) and heated to reflux at 85 ºC for 45 min. The reaction mixture was cooled to 25 ºC, concentrated via rotary evaporation, and purified by flash chromatography using dichloromethane and methanol to obtain the title compound as blue solid (236 mg, 60% yield). $^{1}$H NMR ($CDCl_3$, 500 MHz) δ ppm 8.14-8.08 (m, 1H), 7.88-7.79 (m, 2H), 7.32-7.28 (m, 2H), 7.12 (t, *J* = 7.5 Hz, 1H), 7.01 (d, *J* = 7.5 Hz, 1H), 5.93 (d, *J* = 12.5 Hz, 1H), 3.88 (t, *J* = 8.0 Hz, 2H),

3.38 (s, 3H), 3.36 (s, 3H), 2.35 (t, *J* = 7.0 Hz, 2H), 2.00 (qt, *J* = 7.0 Hz, 2H), 1.65 (s, 6H), 1.49 (s, 9H); $^{13}$C NMR ($CDCl_3$, 125 MHz) δ ppm 171.7, 169.3, 163.8, 162.7, 156.7, 155.7, 152.2, 142.7, 140.1, 128.6, 123.5, 122.0, 121.7, 109.2, 104.5, 100.1, 81.1, 48.2, 42.5, 31.9, 28.32, 28.27, 28.1, 27.6, 21.9; HRMS (positive mode) obsd 494.2649, calcd 494.2650 [$M+H^+$, M = $C_{28}H_{35}N_3O_5$].

**I-BA(4)-NHS**. A solution of compound **8** (236 mg, 0.480 mmol) in dichloromethane (3.0 mL) was treated with trifluoroacetic acid (1.0 mL) and stirred under argon for 2 h. Completion of t-butyl ester deprotection was confirmed by TLC analysis. Then, the reaction mixture was concentrated via rotary evaporation. The resulting residue was mixed with 2,5-dioxopyrrolidin-1-yl 2,2,2-trifluoroacetate (202 mg, 0.960 mmol), 1,2-dichloroethane (5.0 mL) and pyridine (1.0 mL) and stirred under argon for 12 h. The content was mixed with silica gel and evaporated to dryness in a rotary evaporator and purified by flash chromatography using hexane and ethyl acetate. The product was isolated as a dark blue solid (136 mg, 52% yield). $^1$H NMR ($CDCl_3$, 500 MHz) δ 8.11 (d, *J* = 12.5 Hz, 1H), 7.86-7.75 (m, 2H), 7.33-7.28 (m, 2H), 7.12 (t, *J* = 7.5 Hz, 1H), 6.96 (d, *J* = 7.5 Hz, 1H), 6.03 (d, *J* = 12.5 Hz, 1H), 3.96 (t, *J* = 9.0 Hz, 2H), 3.38 (s, 3H), 3.35 (s, 3H), 2.96 (s, 4H), 2.78 (t, *J* = 6.5 Hz, 2H), 2.18 (qt, *J* = 7.5 Hz, 2H), 1.65 (s, 6H); $^{13}$C NMR ($CDCl_3$, 125 MHz) δ ppm 169.2, 168.8, 167.7, 163.7, 162.7, 157.0, 155.5, 152.2, 142.4, 140.0, 128.5, 123.5, 122.1, 108.7, 104.9, 100.3, 48.1, 41.8, 28.4, 28.3, 27.6, 25.4, 21.8; HRMS (positive mode) obsd 535.2186, calcd 535.2188 [$M+H^+$, M = $C_{28}H_{30}N_4O_7$].

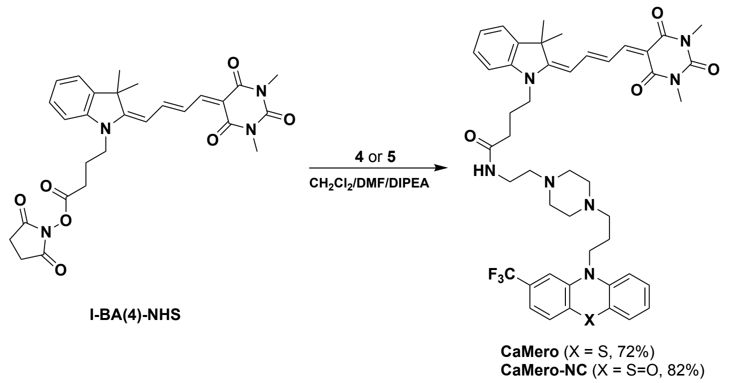


**Scheme S3**. Synthesis of **CaMero** and **CaMero-NC**

**CaMero**. A solution of compound **I-BA(4)-NHS** (28 mg, 0.052 mmol) and 2-(4-(3-(2-(trifluoromethyl)-10H-phenothiazin-10-yl)propyl)piperazin-1-yl)ethan-1-amine **4** (20 mg, 0.046 mmol) in a solvent mixture containing 1,2-dichloroethane (1.0 mL) and DMF (100 µL) was treated with *N,N*-diisopropylethylamine (200 µL) and stirred at 25 ºC under argon for 24 h. The content was mixed with silica gel and evaporated to dryness in a rotary evaporator and purified by flash chromatography using dichloromethane and methanol. The product was isolated as a dark blue solid (28.5 mg, 72% yield). $^1$H NMR ($CDCl_3$, 500 MHz) δ 8.12 (d, *J* = 12.5 Hz, 1H), 7.88-7.80 (m, 2H), 7.32 (t, *J* = 7.5 Hz, 1H), 7.28 (t, *J* = 3.0 Hz, 1H), 7.21-7.11 (m, 6H), 7.05 (s, 1H), 6.96 (t, *J* = 7.5 Hz, 1H), 6.93 (d, *J* = 8.5 Hz, 1H), 6.12 (br s, 1H), 5.95 (d, *J* = 12.5 Hz, 1H), 3.97 (t, *J* = 7.0 Hz, 2H), 3.97 (t, *J* = 6.5 Hz, 2H), 3.40-3.39 (br m, 5 H), 3.37 (s, 3H), 2.40-2.50 (m, 12H), 2.33 (t, *J* = 6.5 Hz, 2H), 2.07 (qt, *J* = 7.0 Hz, 2H), 1.94 (qt, *J* = 7.0 Hz, 2H), 1.65 (s, 6H); $^{13}$C NMR ($CDCl_3$, 125 MHz) δ ppm 171.1, 169.6, 163.8, 162.8, 156.6, 155.8, 152.3, 145.7, 144.3, 142.6, 140.1, 129.9, 129.7, 129.4, 128.5, 127.7, 127.6, 127.5, 125.3, 124.0, 123.6, 123.1, 121.9, 121.6, 119.0 (q, $CF_3$ coupling), 115.9, 111.9 (q, $CF_3$ coupling), 109.6, 104.3, 100.1, 56.4, 55.2, 53.1, 52.8, 48.3, 45.2, 42.8, 35.9, 32.2, 31.6, 28.33, 28.26, 27.6, 25.3, 24.1, 22.7, 22.2; HRMS (positive mode) obsd 856.3829, calcd 856.3827 [$M+H^+$, M = $C_{46}H_{52}F_3N_7O_4S$].

**CaMero-NC**. A solution of compound **I-BA(4)-NHS** (6.7 mg, 0.013 mmol) and 10-(3-(4-(2-aminoethyl)piperazin-1-yl)propyl)-2-(trifluoromethyl)-10H-phenothiazine 5-oxide **5** (9.0 mg, 0.020 mmol) in a solvent mixture containing 1,2-dichloroethane (1.0 mL) and DMF (100 µL) was treated with *N,N*-diisopropylethylamine (100 µL) and stirred at 25 ºC under argon for 5 h. The content was mixed with silica gel and evaporated to dryness in a rotary evaporator and purified by flash chromatography using dichloromethane and methanol. The product was isolated as a dark blue solid (9.3 mg, 82% yield). $^{1}$H NMR ($CDCl_3$, 500 MHz) δ 8.04 (d, $J$ = 12.5 Hz, 1H), 7.98 (d, $J$ = 7.5 Hz, 1H), 7.88 (dd, $J_1$ = 8.0 Hz, $J_2$ = 2.0 Hz, 1H), 7.81-7.70 (m, 2H), 7.65 (s, 1H), 7.59 (dt, $J_1$ = 8.0 Hz, $J_2$ = 2.0 Hz, 1H), 7.50 (d, $J$ = 8.0 Hz, 1H), 7.40 (d, $J$ = 8.0 Hz, 1H), 7.26-7.21 (m, 3H), 7.06-7.02 (m, 2H), 6.26 (br s, 1H), 5.88 (d, $J$ = 12.5 Hz, 1H), 4.37 (td, $J_1$ = 7.0 Hz, $J_2$ = 3.0 Hz, 2H), 3.84 (t, $J$ = 8.0 Hz, 2H), 3.37 (br s, 2H), 3.30 (s, 3H), 3.29 (s, 3H), 2.55-2.42 (m, 12H), 2.25 (t, $J$ = 7.0 Hz, 2H), 2.00-1.97 (m, 4H), 1.57 (s, 6H); $^{13}$C NMR ($CDCl_3$, 125 MHz) δ ppm 171.3, 169.7, 163.8, 162.8, 156.7, 155.9, 152.3, 142.6, 140.1, 138.9, 138.1, 133.1, 132.1, 131.48.3, 4, 123.6, 122.8, 121.9, 121.6, 118.0, 116.4, 113.3, 109.6, 104.2, 100.2, 56.5, 54.1, 53.4, 52.6, 45.2, 42.8, 35.6, 32.2, 29.7, 28.33, 28.27, 27.6, 23.8, 22.7, 22.2; HRMS (positive mode) obsd 872.3766, calcd 872.3776 [$M+H^+$, M = $C_{46}H_{52}F_3N_7O_5S$].

**Photophysical characterization of biosensor**. UV-visible spectra were obtained with a Hewlett-Packard 8453 diode array spectrophotometer or Agilent Cary 60 UV-Vis spectrophotometer. Emission and excitation spectra were obtained using a Spex Fluorolog 2 spectrofluorometer at 25 °C. Instrument-specific correction factors provided by the instrument manufacturer were applied to the raw fluorescence data. The final data were reported in the corrected signal/corrected reference mode.

In the experiment to study the response of biosensor to calmodulin, CaMero (1 µM) was added to an aqueous Tris buffer containing $Ca^{2+}$ (Free calcium concentration = 10 µM). NaCl was added to reach an ionic strength of 150 mM. The solution was excited at 545 nm and emission was collected from 560 to 650 nm. Then, calmodulin (1 µM), EGTA ((Free calcium concentration = 0.2 µM)) and $Ca^{2+}$ (Free calcium concentration = 100 µM) were sequentially added, and the emissions were collected for each event. Emission was normalized for CaMero concentration, and intensity in the wavelength window 589 – 625 nm was plotted for each event.

### REFERENCES


1 MacNevin, C. J. *et al.* Membrane-Permeant, Environment-Sensitive Dyes Generate Biosensors within Living Cells. *Journal of the American Chemical Society* **141**, 7275–7282 (2019). https://doi.org/10.1021/jacs.8b09841
